# Single-molecule imaging with cell-derived nanovesicles reveals early binding dynamics at a cyclic nucleotide-gated ion channel

**DOI:** 10.1101/2021.07.05.451181

**Authors:** Vishal R. Patel, Arturo M. Salinas, Darong Qi, Shipra Gupta, David J. Sidote, Marcel P. Goldschen-Ohm

## Abstract

Ligand binding to membrane proteins is critical for many biological signaling processes. However, individual binding events are rarely directly observed, and their asynchronous dynamics are occluded in ensemble-averaged measures. For membrane proteins, single-molecule approaches that resolve these dynamics are challenged by dysfunction in nonnative lipid environments, lack of access to intracellular sites, and costly sample preparation. Here, we introduce an approach combining cell-derived nanovesicles, microfluidics, and single-molecule fluorescence colocalization microscopy to track individual binding events at a cyclic nucleotide-gated TAX-4 ion channel critical for sensory transduction. Our observations reveal dynamics of both nucleotide binding and a subsequent conformational change likely preceding pore opening. We further show that binding of the second ligand in the tetrameric channel is less cooperative than previously estimated from ensemble-averaged binding measures. This approach is broadly applicable to studies of binding dynamics for proteins with extracellular or intracellular domains in native cell membrane.

## Introduction

Ligand binding to specific recognition sites in membrane receptors is crucial for cellular signaling and pharmacological treatment of its dysfunction. Intrinsic membrane proteins make up ∼30% of the protein encoding genome and are therapeutic targets for ∼70% of available drugs^1–3^. Technological advances have led to an increase in near atomic resolution structures of membrane receptors complexed with ligands/drugs. These structures provide static snapshots primarily of endpoints for ligand-activation mechanisms. However, the transient intermediate events connecting these snapshots often remain unclear. For oligomeric proteins that bind ligand/drug at multiple active sites, these intermediates define the sequence of binding events and the nature of cooperative interactions amongst sites (i.e. occupation of one site influencing binding at another) that govern the concentration-dependence of ligand-induced behavior. Methods that inform on these processes are important for understanding mechanisms of ligand-activation and aiding the rational design of novel therapies targeting membrane receptors. However, structures and ensemble-averaged measures of distinct partially liganded intermediates are difficult to resolve due to averaging over transient asynchronous events and mixtures of heterogeneous bound states.

Single-molecule (SM) approaches are ideal for resolving both heterogeneous states and asynchronous dynamics. However, their application to studies of ligand binding in membrane proteins is challenged by costly sample preparation, dysfunction in nonnative lipid or detergent environments^4–6^, lack of solution access to intracellular sites, and nonspecific dye adsorption to imaging surfaces. Here, we overcome these challenges by combining cell-derived nanovesicles^7–9^, microfluidics, micro-mirror total internal reflection fluorescence (mmTIRF)^10^, and colocalization SM spectroscopy^11^ to optically track binding and unbinding of individual fluorescently-tagged ligands at single membrane proteins. Imaging a ∼100 μm × 100 μm field of view much larger than a confocal spot enables high-throughput data acquisition of up to hundreds of molecules simultaneously.

Cell-derived nano-scale vesicles are an attractive approach for isolating full-length membrane proteins in their native lipid environment and provide access to extracellular and intracellular sites due to the stochastic nature of vesicle formation. SM imaging of individual vesicles has been used to study nicotinic acetylcholine receptor (nAChR) stoichiometry^8^ and mechanisms of membrane curvature sensing^12^. In contrast, live cell-based approaches offer a native membrane environment^13–15^ but are challenged by cell autofluorescence, diffusion within the membrane, high local protein concentrations that limit SM resolution^16–18^, and lack of access to intracellular domains. Isolating proteins with patch pipets allows simultaneous optical and electrical recording^19^, but severely limits throughput to one molecule at a time. Alternative cell-free approaches such as solubilization in detergent or lipid synthetics can leave the protein locked into conformations that constrain normal function^4–6^ or require stabilizing mutations that may impose unnatural constraints on protein structure and function^20, 21^.

As an exemplar system, we use our approach to resolve the binding dynamics of a fluorescent cyclic nucleotide analog (fcGMP^22, 23^) to individual cyclic nucleotide-gated (CNG) TAX-4 ion channels from *C. elegans* in cell-derived nanovesicles. CNG channels are critical for visual and olfactory transduction^24^. They are tetramers comprised of four subunits surrounding a central ion conducting pore^25^. Binding of cGMP or cAMP to intracellular cyclic nucleotide binding domains (CNBDs), one per subunit, initiates opening of the channel’s cation conducting pore, thereby converting changes in cyclic nucleotide level to changes in membrane potential^24^. This allows photoreceptors in the visual system to respond to changes in light level via light-induced hydrolysis of cGMP, and olfactory receptor neurons to respond to odorant-induced synthesis of cAMP^24, 26^. Mutations in CNG channels have been linked to progressive vison loss and color vision abnormalities such as macular degeneration^27–31^ as well as olfactory disorders such as isolated congenital anosmia^32^. Recently, a clinical trial of subretinal *CNGA3* gene therapy in individuals with complete achromatopsia resulted in significantly improved visual acuity and contrast sensitivity, thus validating CNG channels as a promising therapeutic target^33^.

Cryo-EM structural models of TAX-4^34, 35^, human rod CNGA1 channels^36^, and prokaryotic homologs^37, 38^ in both unliganded and cGMP/cAMP-bound conformations provide important static snapshots of endpoints during ligand activation. These structures as well as others from prokaryotic homologs^37, 38^ are complimented by functional studies of channel currents and measures of ligand-dependent conformational changes using approaches such as fluorescence spectroscopy^39^, electron paramagnetic resonance^40^, and high-speed atomic force microscopy^41^. However, these studies do not by themselves reveal the sequence of events connecting the observed structural endpoints, which requires temporal resolution of distinct bound states that have previously not been directly observed.

Our observations provide a first look at individual CNBD dynamics and initial binding cooperativity in a full-length CNG channel embedded in native cell membrane. These results reveal similarity to individual site dynamics with structurally similar CNBDs from HCN channels and constrain plausible models of binding cooperativity. Our combined approach has broad application to other membrane receptors and complements structural information with dynamics for transient states whose structures are not easily resolved.

## Results

### Immobilization of cell-derived nanovesicles for single-molecule imaging

For imaging and immobilization, we fused the enhanced green fluorescent protein (EGFP) to the cytosolic N-terminus of the CNG channel TAX-4 (GFP-TAX-4). Cell-derived nanovesicles containing GFP-TAX-4 were generated as described for a study of nicotinic acetylcholine receptor stoichiometry^7–9, 42^ (Fig. 1). Briefly, HEK-293T cells transfected with GFP-TAX-4 were disrupted using nitrogen cavitation, resulting in the spontaneous formation of cell membrane vesicles with diameters around 200 nm^9, 42^ (Supplementary Fig. 1). The vesicles were further separated by gradient ultracentrifugation into fractions comprised primarily of either plasma membrane (PM) or endoplasmic reticulum (ER) membrane (Supplementary Fig. 2). The PM vesicle fraction was applied to a microfluidic chamber on a passivated glass coverslip with a GFP-nanobody bait protein sparsely deposited on its surface for on-chip purification and immobilization of GFP-TAX-4^43, 44^. Due to the stochastic nature of vesicle assembly, a mixture of vesicles with either extracellular or intracellular leaflets exposed to the bath solution are obtained^45^. Vesicles containing GFP-TAX-4 oriented with both GFP and the intracellular CNBDs outside the vesicle are immobilized upon binding to the GFP-nanobody on the chip surface (Fig. 1), whereas vesicles without GFP-TAX-4 or with GFP and CNBDs oriented towards the inside of the vesicle are washed away upon rinsing with buffer (Supplementary Fig. 3a). To verify the applicability of this technique to proteins with more complex stoichiometries and extracellular binding domains such as reported for nACHRs^8^, we performed the same nanovesicle preparation and immobilization procedure with hetero-pentameric GABA_A_ receptors comprised of *α*_1_, *β*_2_ and *γ*_2L_ subunits (Supplementary Fig. 3b).

**Fig. 1.**
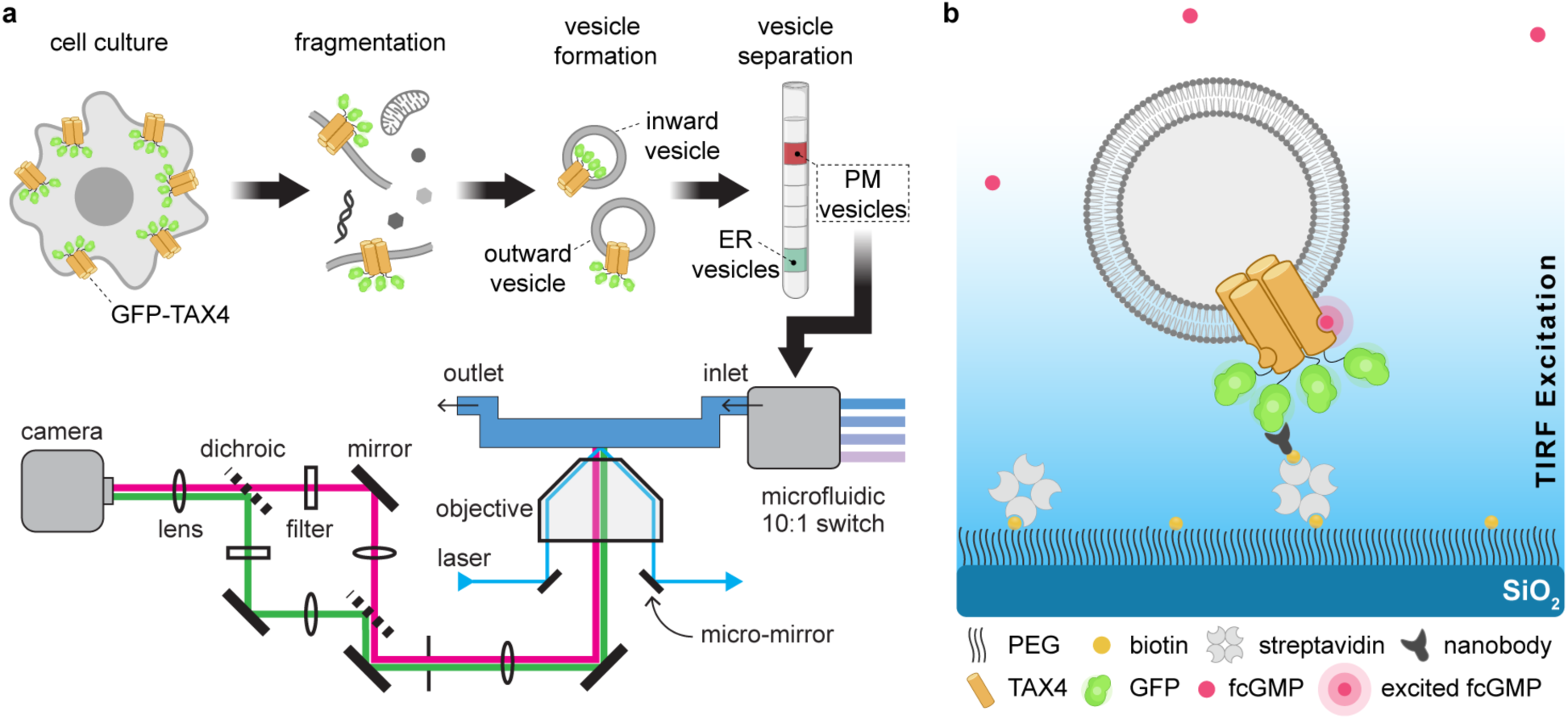
Immobilization of cell-derived nanovesicles for single-molecule imaging of membrane proteins. **a** Schematic illustrating both sample preparation and the imaging setup. Briefly, cells expressing the membrane protein of interest (e.g. GFP-TAX-4) are fragmented using N_2_ cavitation to spontaneously form nanoscale vesicles, some of which contain the protein of interest in a mixture of inward and outward facing orientations. Vesicles comprised of membrane from either the plasma membrane (PM) or endoplasmic reticulum (ER) are further separated by gradient ultracentrifugation. The fraction of PM vesicles is applied to a sample chamber for immobilization and imaging with micro-mirror total internal reflection (mmTIRF) microscopy. **b** Illustration of an individual immobilized vesicle in the sample chamber. The chamber consists of a glass coverslip coated with a layer of PEG doped with PEG-biotin to which a biotinylated anti-GFP nanobody (bait protein) is attached via streptavidin. Vesicles containing GFP-TAX-4 oriented such that GFP is exposed to the extravesicular solution are immobilized at nanobody locations on the optical surface. TIRF excitation, indicated by the blue gradient, ensures that bulk freely diffusing fluorescent ligand (e.g. fcGMP) above the vesicle layer are not appreciably excited. Fluorescence from the vesicle layer is imaged on an EMCCD as depicted to the left. Note that the vesicle is not drawn to scale as it would on average have a diameter about 20-fold larger than GFP-TAX-4.

### Resolving individual binding events at TAX-4 CNG channels

For optical detection of ligand binding events the sample chamber was continuously perfused with a fluorescent cGMP conjugate (fcGMP) previously shown to activate CNG channels with similar efficacy and affinity to cGMP^22^. Importantly, TIRF limits excitation laser power to an evanescent field within ∼100-200 nm of the optical surface, encompassing primarily the layer of immobilized vesicles only. This is critical, as illumination of the vast majority of freely diffusing fcGMP above the vesicle layer would result in background fluorescence precluding resolution of individual fcGMP molecules. Because the time for free fcGMP to diffuse through a diffraction-limited spot (∼1 ms) is appreciably shorter than the time resolution of our recordings (50 ms per frame), we did not resolve diffusion of fcGMP to or from TAX-4, but only observed increased fluorescence when fcGMP remained at a spot for a time period comparable to or longer than the frame duration, as when bound to TAX-4.

Colocalized spots with both EGFP and fcGMP fluorescence were identified as immobilized vesicles containing GFP-TAX-4 with functional binding domains (Fig. 2). To determine whether colocalized binding events represent specific binding to TAX-4 rather than nonspecific adsorption to the surface or lipids, we tested the ability of nonfluorescent cGMP to outcompete fcGMP for its binding site. Using a microfluidic pump and switch we perfused a mixture of fcGMP and an excess of nonfluorescent cGMP to outcompete fcGMP binding at the same molecules previously imaged in fcGMP alone. Colocalization of EGFP and fcGMP fluorescence, consistent with specific binding to immobilized GFP-TAX-4 receptors, was largely abolished by competition with nonfluorescent cGMP and recovered upon reapplication of fcGMP alone (Fig. 2). This suggests that reversible binding at these locations reflects fcGMP specific association with TAX-4 CNBDs.

**Fig. 2.**
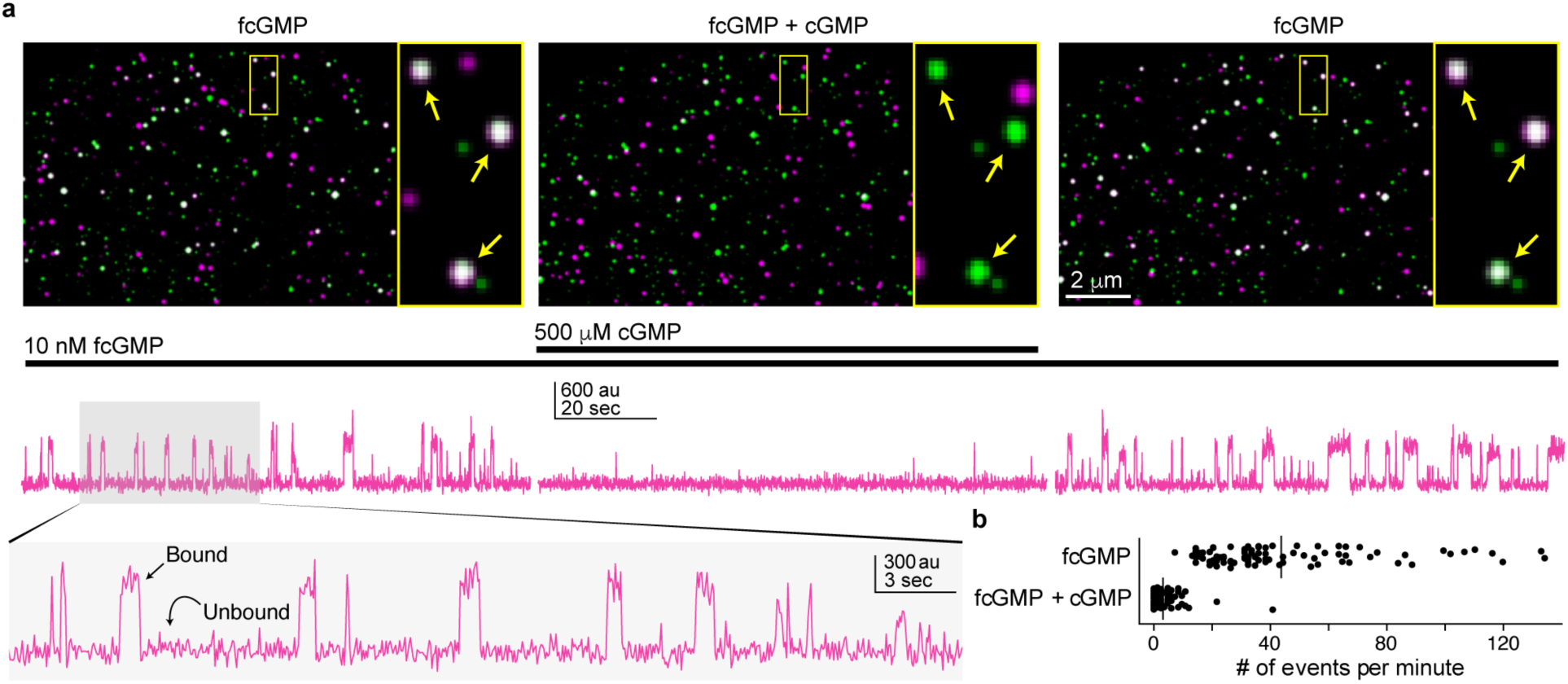
Single-molecule imaging of fcGMP binding to GFP-TAX-4 in cell-derived nanovesicles. **a** Time-averaged fluorescence for GFP (green) overlaid with fcGMP (magenta). Colocalized GFP and fcGMP signals appear white. From left to right depicts sequential epochs for the same field of view showing binding in 10 nM fcGMP, block of specific binding by coapplication with an excess of 500 μM non-fluorescent cGMP, and recovery upon removal of cGMP. The yellow box is expanded to the right of each image and arrows indicate locations exhibiting specific fcGMP binding to GFP-TAX-4. Below each image are time series for fcGMP fluorescence at a single colocalized spot during each of the epochs in the corresponding images above. Transient increases in fluorescence reflect individual fcGMP binding events in the vesicle layer at the optical surface. The shaded region is expanded below. **b** The frequency of binding events in 10 nM fcGMP is greatly diminished in the presence of 500 μM competing non-fluorescent cGMP.

We also observed fcGMP fluorescence at spots devoid of EGFP signal. The similar frequency of these noncolocalized spots upon competition with cGMP suggests that they reflect nonspecific adsorption of fcGMP (Fig. 2). To characterize noncolocalized signals, we compared the mean intensity and standard deviation of individual events to those at colocalized spots. Noncolocalized fcGMP events tend to have lower intensities and higher variance than colocalized events (Supplementary Fig. 4a). However, we occasionally observed noncolocalized high intensity events that we interpret as adsorption of fcGMP to the coverslip where the TIRF excitation field is most intense (Supplementary Fig. 4b-c). In this case, the lower intensity noncolocalized events are likely to reflect adsorption to aggregates in the surface layer. The high intensity events were generally much longer lived than the vast majority of binding events, suggesting that termination of most colocalized binding events can be attributed to unbinding rather than fluorophore bleaching. Noncolocalized EGFP spots were also observed, which we hypothesize reflect vesicles where surface interactions occlude solution access to CNBDs or otherwise render them nonfunctional. To verify that colocalized fcGMP fluorescence signals were not attributable to nonspecific interactions with nanovesicle lipids, we performed the same fcGMP binding experiment with immobilized nanovesicles containing either a TRPV1 channel fused to an intracellular GFP or GABA_A_ receptors that do not bind cGMP. We did not observe any appreciable colocalization in the absence of GFP-TAX-4 (Supplementary Figs. 5-6). Taken together, these observations strongly suggest that colocalized EGFP and fcGMP florescence spots reflect specific binding of individual fcGMP molecules to the CNBDs of immobilized GFP-TAX-4 channels.

To restrict our analysis to colocalized spots containing single CNG channels we estimated the number of GFP-TAX-4 subunits in each diffraction-limited spot by counting the number of bleach steps in the EGFP fluorescence time series (Fig. 3a). Given that TAX-4 channels are tetramers, we excluded from the analysis all spots with more than four bleach steps. The distribution of bleach steps for the remaining spots was well described by a binomial distribution for four sites with an estimated probability of observing each EGFP bleaching event of 0.82, consistent with reports for this probability (Fig. 3b)^8, 14^. These spots were considered as having a high probability of containing single tetrameric GFP-TAX-4 channels.

**Fig. 3.**
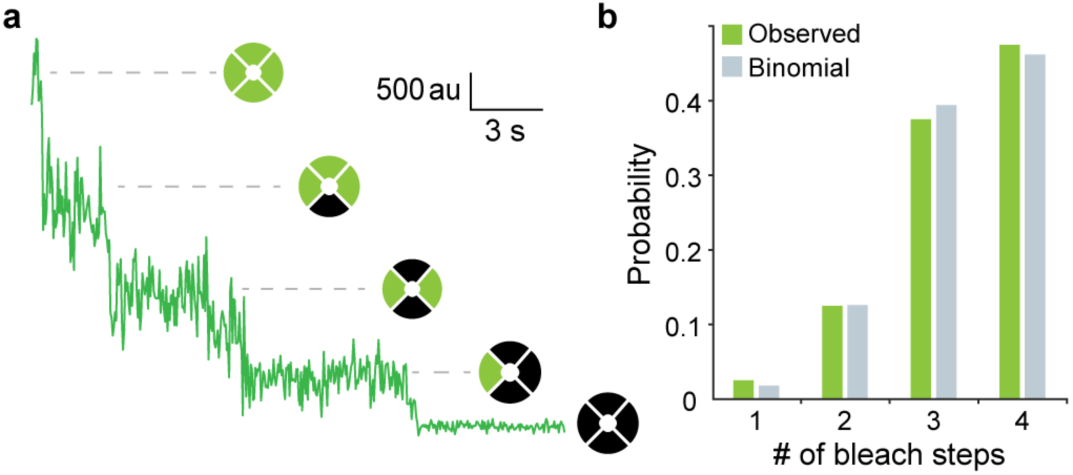
Number of GFP-TAX-4 subunits per vesicle assessed by photobleaching of GFP. **a** Stepwise photobleaching of GFP fluorescence from a single diffraction-limited spot. **b** The number of observed bleach steps for colocalized spots included in the analysis. The good fit to a binomial distribution with four sites suggests that most spots contain a single tetrameric channel where bleaching of each GFP is observed with a probability of 0.82 and 95% CI [0.74, 0.89].

### TAX-4 CNBDs undergo a conformational change following binding

We measured a total of ∼60 hours of SM binding dynamics across a range of fcGMP concentrations from 10-200 nM. Time series for fcGMP binding at these spots show a clear dependence on concentration as expected for binding events (Fig. 4a). At low concentrations of 10-30 nM fcGMP we almost exclusively observed single isolated binding events. With increasing concentrations from 60-200 nM fcGMP we observed an increasing frequency of simultaneous binding of multiple fcGMP at individual molecules (i.e. stacked fluorescence steps). We did not explore higher fcGMP concentrations necessary to saturate the binding sites due to the challenge of increased background fluorescence from freely diffusing ligand.

**Fig. 4.**
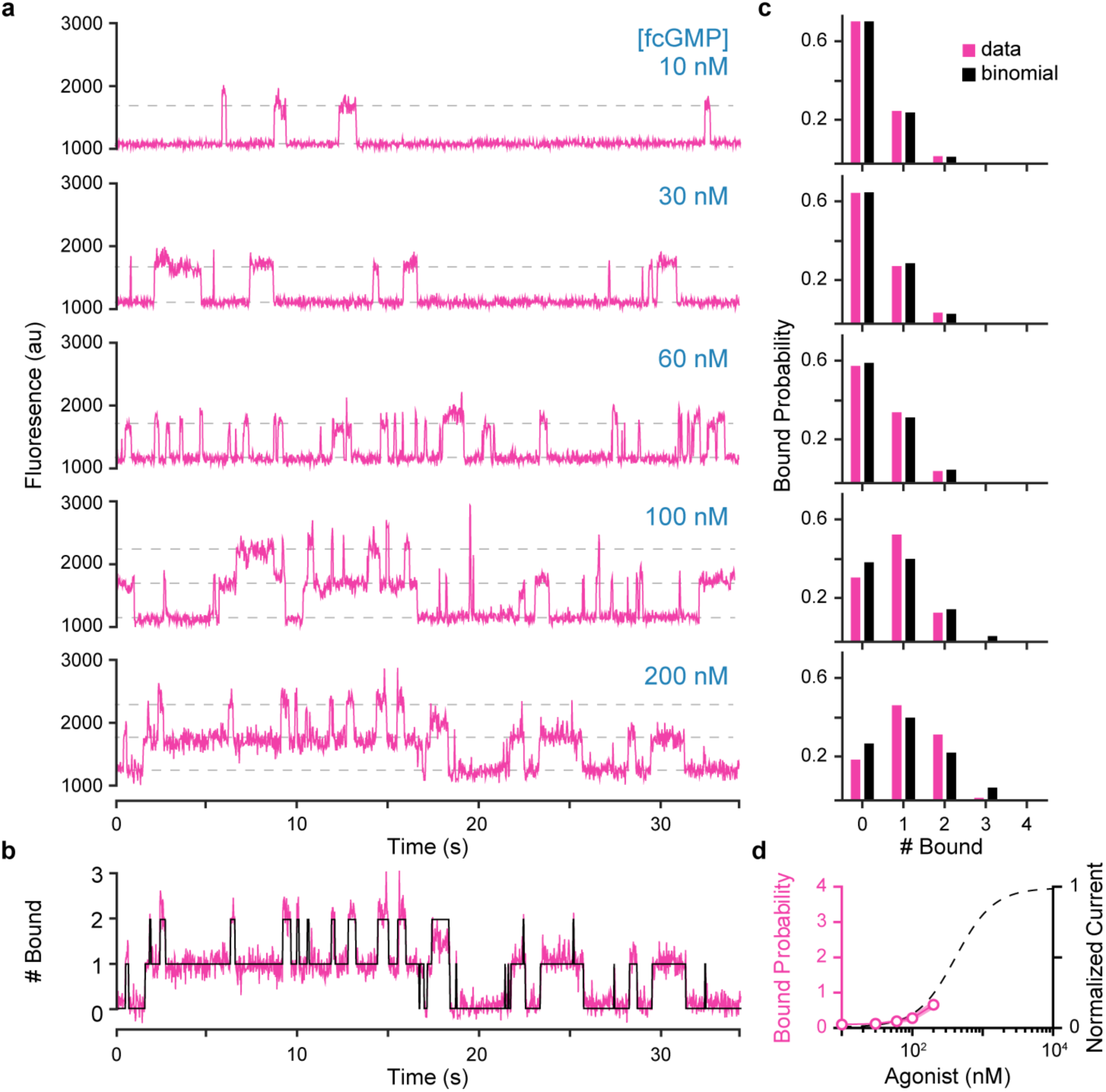
Concentration-dependent fcGMP binding. **a** Time series for fcGMP binding to single TAX-4 containing nanovesicles at increasing concentrations of fcGMP. Horizontal dashed lines indicate approximate fluorescence levels reflecting stepwise binding of one or two fcGMP molecules. **b** Idealization (black) of fluorescence (magenta) time series for the number of bound fcGMP at an individual molecule in 200 nM fcGMP (see Methods). **c** Bound probability distributions for all tested concentrations and fits to a binomial distribution assuming four identical and independent sites. Bound probability [95% CI] per site from the binomial fits: 10 nM = 0.08 [0.079, 0.081], 30 nM = 0.10 [0.09, 0.11], 60 nM = 0.12 [0.10, 0.14], 100 nM = 0.21 [0.20, 0.22], 200 nM = 0.26 [0.24, 0.28]. **d** Average fcGMP bound probability across all molecules and normalized ionic current^46^ as a function of fcGMP or cGMP concentration, respectively. Shaded area represents SEM (partially hidden by the line width). Current values are for fits of the Hill equation to inside-out patch clamp recordings in Komatsu et al. (1999)^46^.

Fluorescence time series for fcGMP binding were idealized to obtain the number of bound fcGMP at each time point (Fig. 4b; Supplementary Figs. 7-8; see Methods). The average bound probability across molecules exhibited a dependence on fcGMP concentration similar to observations of the cGMP concentration dependence of channel current^46^ (Fig. 4d), further supporting the idea that our observations reflect binding events associated with activation of TAX-4. Furthermore, bound lifetimes were independent of concentration, whereas unbound lifetimes monotonically decreased with increasing fcGMP concentration as qualitatively expected for a binding reaction (Fig. 5a,c). For periods with simultaneous occupation of multiple sites, we estimated the bound lifetime at each site by randomly assigning each unbinding event to one of the bound molecules. Repeating this random assignment led to nearly identical distributions, indicating that randomization did not severely distort the distributions. We assessed the ability of the idealization procedure to resolve individual events by applying it to simulated fluorescence binding data with lifetimes and noise drawn from the experimental observations (Supplementary Fig. 7). Comparison of the simulated and idealized event records indicates overall very good detection of singly and doubly liganded events except for brief single frame events in doubly liganded states which were missed approximately one third of the time (Supplementary Fig. 8; see Methods).

**Fig. 5.**
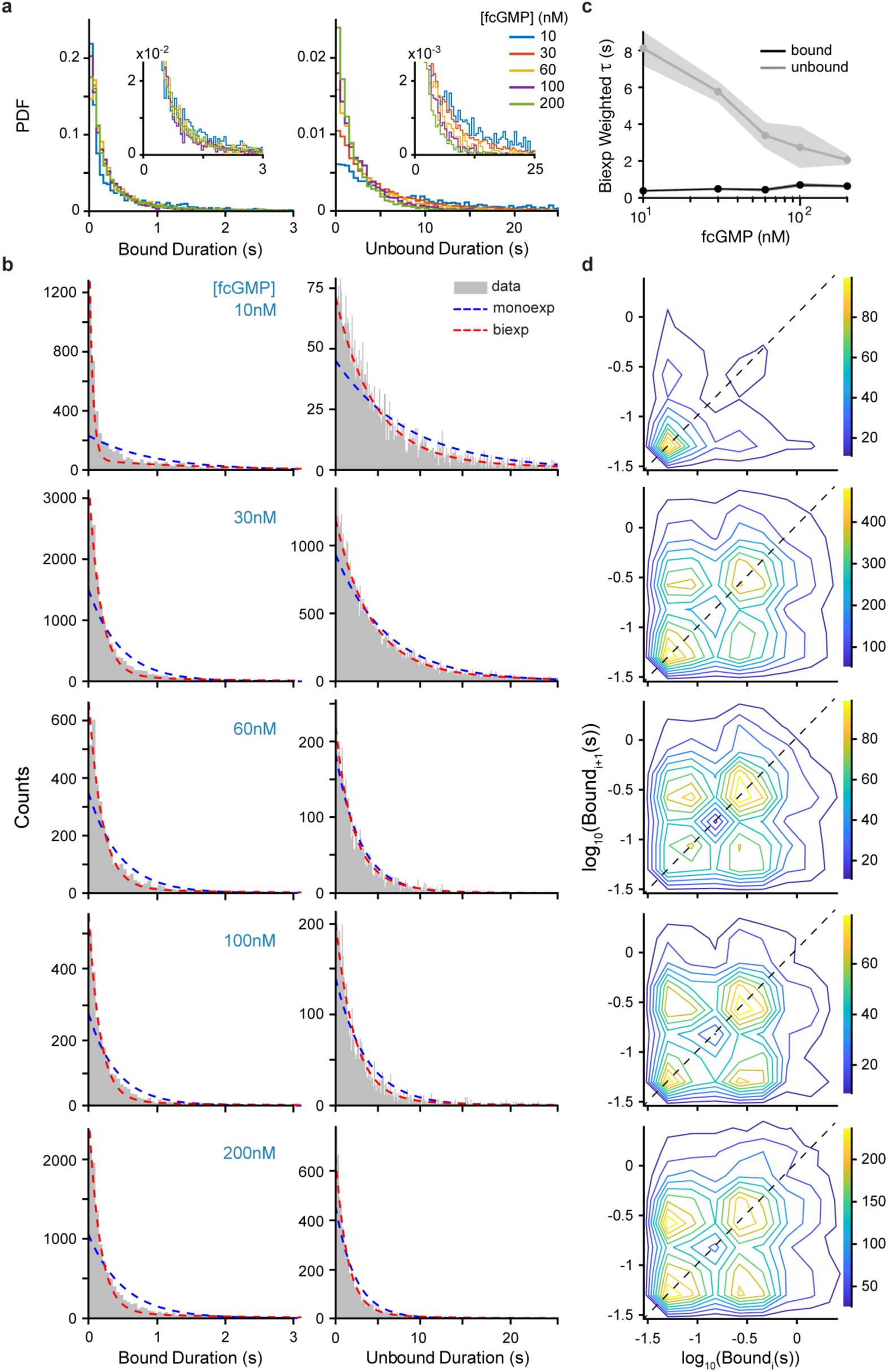
Bound and unbound dwell time distributions and correlations. **a** Bound and unbound fcGMP dwell time distributions across all molecules from idealized records (see Methods). **b** Dwell time distributions for data (gray) overlaid with mono-(blue dashed) and bi-exponential (red dashed) maximum likelihood fits (see Supplementary Table 1). **c** Weighted time constants from biexponential fits of bound (black) and unbound (gray) dwell times as a function of fcGMP concentration with associated 95% confidence intervals (shaded area). **d** Correlation between the duration of sequential singly bound events *i* and *i*+1 within individual molecules. The concentration of fcGMP is the same as for the dwell time distributions to the left. Color bar denotes number of events. If short and long events arise from distinct populations of molecules, we would expect to observe clusters of events primarily along the dashed diagonal, whereas events on the off-diagonal represent sequential short and long bound durations within individual molecules.

Bound lifetimes at all tested fcGMP concentrations were poorly described by a single exponential distribution indicative of unbinding from a singular bound state. Instead, at least two exponential components were needed to account for our observations (Fig. 5b; Supplementary Table 1), suggesting that the CNBDs adopt multiple bound conformations. To determine whether these different conformations arise from two distinct populations of molecules or reflect conformational exchange within a single population of molecules, we examined correlations between the lifetimes of successive binding events at individual molecules (Fig. 5d). If shorter and longer bound lifetimes were attributable to distinct populations of molecules, we would expect little correlation between short and long bound events within individual molecules. In this case we would expect to see primarily either short- or long-lived binding events at an individual molecule (i.e. lower left or upper right along the dashed diagonal line in Fig. 5d), and relatively fewer pairs of sequential short/long or long/short events (i.e. the off-diagonal in Fig. 5d). In contrast, we observe both short and long binding events at individual channels with high probability, suggesting that individual CNBDs exchange between at least two bound conformations. Unbound lifetime distributions were also better described by two exponentials, suggesting that unliganded CNBDs also adopt at least two conformations. However, one of the two exponential components accounted for the majority of the unbound lifetime distributions, consistent with high frequency occurrence of a single unliganded conformation.

To explore the dynamics of this process at individual CNBDs we evaluated a series of hidden Markov models (HMMs) ranging from the simplest possible two state binding mechanism to models with up to two bound and unbound states (Supplementary Fig. 9). We restricted analysis of these models to isolated binding events by removing periods with two or more bound ligands, thereby splitting those time series into segments comprised only of singly bound events. Models were globally optimized in QuB^47, 48^ for all molecules and concentrations and ranked according to their relative Bayesian Information Criterion (*Δ*BIC = BIC – BIC_best model_) scores (smaller is better) (Supplementary Fig. 9; Supplementary Table 2, see Methods). As expected, given the observed lifetime distributions, models with two bound states were preferred (smaller *Δ*BIC) over those with a single bound state. Addition of a second unbound state also reduced *Δ*BIC, although to a lesser extent than addition of a second bound state. Both models with cyclic reaction schemes (M1.D and M1.F) resulted in much slower rate constants for one of the transition pairs, suggesting that a full cycle is traversed relatively infrequently. The best ranked model M1.F predicts that binding and unbinding following the conformational change is an order of magnitude slower than that preceding it (Supplementary Table 2). Furthermore, the *Δ*BIC score for model M1.E which disallows binding/unbinding after the conformational change was almost equivalent to that of M1.F. Taken together, these observations suggest that binding following the conformational change is at least inhibited, and thus we favor a simplified sequential model that lacks these transitions. Our favored model M1.E (Fig. 6a; Supplementary Fig. 9) has the same form as the model describing dynamics at individual isolated CNBDs from HCN2 channels^49^. Although M1.E predicts that the conformational change can occur in the absence of ligand, rates of exchange are much slower than those for ligand-bound sites (Supplementary Table 2). Thus, the relatively simpler model M1.B with only a single unbound state may also provide a reasonable simplified description of the data. In either case, the two best ranked models M1.E and M1.F as well as the simplified model M1.B all indicate that the CNBD undergoes a conformational change following binding, which we hypothesize involves capping of the binding pocket by the C helix as we previously proposed for HCN2 CNBDs^49^ (Fig. 6a).

**Fig. 6.**
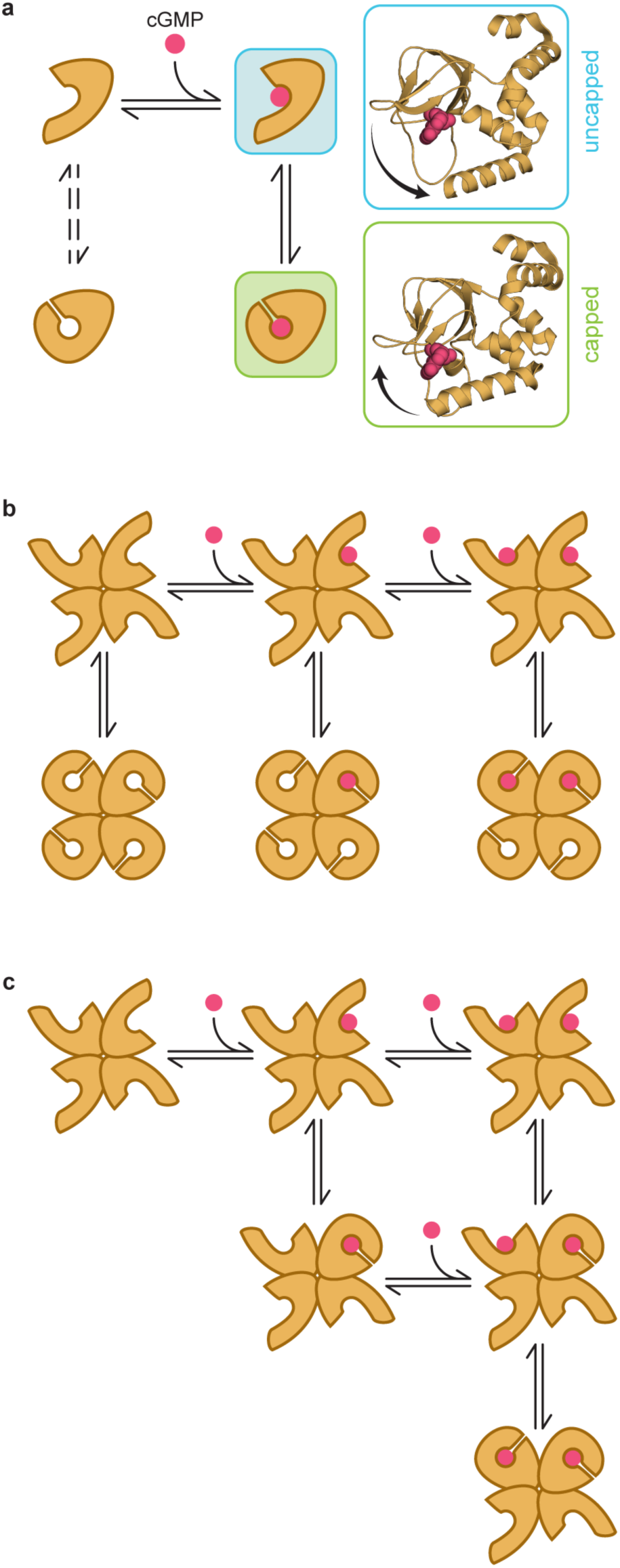
Models for the first and second binding steps. **a** The preferred model for binding dynamics at individual CNBDs. This model depicts both a ligand association step (horizontal transition) as well as a conformational change of the CNBD in both unliganded and liganded states (vertical transitions). The dashed arrows indicate transitions that have a relatively smaller contribution to the observed dynamics. Postulated structures for the ligand-bound conformations are shown to the right. They are based on cryo-EM structures of TAX-4 in cGMP-bound (green) or unliganded (blue) conformations^34, 35^, where cGMP has been added to the unliganded structure in the position that it is found in the bound structure, and arrows denote the observed movement of the C helix between the two structures. Capping of the binding site by the C helix limits binding and unbinding, similar to dynamics observed at isolated CNBDs from HCN2 channels^49^. See Supplementary Fig. 9 and Supplementary Table 2 for all explored models of individual CNBDs and their optimized rate constants. **b-c** Two potential models depicting binding of the first two ligands and either a global conformational change of all four CNBDs in each ligation state (b) or individual conformational changes of each ligand-bound CNBD (c). We were unable to distinguish between these mechanisms. See Supplementary Fig. 10 and Supplementary Table 3 for all explored models of the first two binding steps and their optimized rate constants.

### The first two binding steps

A fundamental limitation to fluorescence imaging is that background fluorescence and noise increase with increasing concentration of fluorescent dye. At micromolar concentrations of fcGMP, this background noise challenges identification of individual binding events. Thus, we limited our observations to concentrations that did not saturate the four binding sites in individual channels. Nonetheless, we were still able to resolve simultaneous ligand binding events as distinguished by multiple stepwise increases in fluorescence. At the highest concentration tested (200 nM) most events (99%) reflect 0, 1, or 2 bound ligands. We therefore restricted our analysis to only the first two binding steps by removing periods with more than two bound ligands from the binding time series, thereby splitting those records into segments. We explored the ability of several two-site HMM models to explain the observed binding dynamics across all molecules and concentrations and ranked them according to their relative *Δ*BIC scores (Supplementary Fig. 10, Supplementary Table 3, see Methods). Note that subscripts on the model labels indicate different sets of constraints as described in Supplementary Fig. 10 and Supplementary Table 3.

The simplest models with two sequential binding steps only (M2.A_i/c_) are much less preferred than models with an additional conformational change in bound and/or unbound ligation states (M2.C_i/c_, M2.D, M2.E_ii/ci/ic_), irrespective of whether or not the binding sites were constrained to be independent (M2.A_i_) or allowed to be cooperative (M2.A_c_). We also examined a model with two distinct doubly liganded states (M2.B; e.g. adjacent and diagonally opposed occupied sites as depicted in Supplementary Fig. 10a) to assess whether distinct patterns of site occupation in the channel tetramer could explain our observations without requiring postulation of a conformational change independent of ligand association/dissociation. Model M2.B had a slightly better *Δ*BIC score than models M2.A_i/c_, but the score for models including a non-binding conformational change were much better (Supplementary Fig. 10c). Together, these results suggests that multiple bound conformations for distinct ligation states is an important feature of cGMP association with TAX-4, consistent with our observations for dynamics at single sites.

To explore the nature of the postulated conformational exchange within each ligation condition we evaluated several models that treat this process as either a global exchange of both CNBDs (M2.C, M2.D) or separate exchanges within each individual CNBD (M2.E). To simplify the models, we did not consider the combination of distinct di-liganded orientations (e.g. M2.B) and conformational changes separate from ligand association/dissociation. Instead, we focused on the conformational change whose importance is suggested by both single site dynamics and the preference for models M2.C-E over M2.A-B. Simulated bound and unbound dwell time distributions for each optimized model describe our observed distributions well for all models that contain multiple bound conformations but differ substantially for models that lack this feature (Supplementary Fig. 11). The best ranked models were M2.C_i/c_ and M2.D. Model M2.D describes a Monod-Wyman-Changeux (MWC) model where ligand affinity depends on the global conformation and the equilibrium between conformations is influenced by the number of bound ligands. M2.C_i/c_ differ from M2.D in that the binding sites are assumed to be accessible in only one conformation, similar to that observed for single sites. Although similarly ranked, optimized parameters for M2.D imply that each successive binding step slows the dynamics of the conformational exchange by roughly an order of magnitude and that association kinetics in the alternate conformation are likewise slowed. To explore whether a different set of parameters for M2.D might be equally likely we constrained binding following the conformational change to be no slower than binding preceding it, in which case the likelihood dropped such that it ranked worse than all other models with the exception of M2.A_i/c_ (not shown). Given that both single site dynamics and M2.D suggest that binding is less favorable following the conformational change, we favor the simpler scheme M2.C where binding/unbinding is disallowed after the conformational change (Fig. 6b). M2.C further predicts that successive binding events bias the conformational equilibrium as qualitatively expected if the conformational exchange is part of the activation process. Scheme M2.C is also consistent with our hypothesis that the conformational change involves capping of the binding pocket by the C helix^49^ (Fig. 6a), and further suggests that binding at one site induces capping of both CNBDs (possibly all four).

Scheme M2.E (Fig. 6c) differs from M2.C and M2.D in that rather than a global conformational exchange of both CNBDs, each CNBD can individually exchange between two bound conformations. Although the likelihood of models M2.E_ii/ci/ic_ are slightly lower than that of models describing a global conformational change (i.e. *Δ*BIC scores are slightly higher), it is not clear that this mechanism can be ruled out. Therefore, whether conformational exchange occurs separately for each CNBD or involves a global rearrangement of both/all CNBDs is uncertain. Additional binding data for the third and fourth step will likely be needed to distinguish these possibilities.

### Binding cooperativity

The distribution of observed average probabilities for 0-4 bound ligands shifted to higher occupation states with increasing concentration. Across concentrations, the distribution was qualitatively described by a binomial distribution as expected for binding at four identical and independent sites (Fig. 4c). However, at higher concentrations the binomial fits tended to underestimate occupation probability, which could reflect cooperative interactions between binding sites (Fig. 4c). We compared models where the first and second binding steps were constrained to be independent (M2.C_i_, M2.E_ii/ic_) or allowed to be cooperative (M2.C_c_, M2.E_ci_) (see constraints in Supplementary Table 3). In all cases, removing the constraint for independent association at each site led to models predicting positive cooperativity for the binding equilibrium of the second ligand (Supplementary Table 3). However, cooperative effects on equilibrium constants were minor, with factors in the range 2.3-3.7. Also, the improvement in *Δ*BIC score from allowing cooperative binding was uniformly minor in comparison to that for addition of a second bound conformation, global or otherwise. We also examined a model where the per-subunit conformational change was allowed to be cooperative (M2.E_ic_) with similar results.

To further explore cooperative binding and to verify our ability to resolve such effects, we compared our experimental data to simulations of independent binding sites (see Methods). For independent sites, the dwell time distribution for events starting upon binding the second ligand and ending upon unbinding of either of the bound ligands should be truncated as compared to the distribution of dwell times for singly bound receptors. This is because statistically the rate of unbinding at either of two identical and independent sites should be twice that of a single site, thereby limiting the frequency of long-lived doubly bound durations compared to singly bound durations. Indeed, simulations of independent sites do predict fewer longer-lived periods with two bound ligands as opposed to one bound ligand, which we can resolve with our idealization procedure after adding noise analogous to our experimental recordings (Supplementary Fig. 12a,b). In contrast, we do not observe a similar reduction in the frequency of longer-lived doubly bound events in our experimental fcGMP binding series (Supplementary Fig. 12c). It is difficult to rule out that such an effect might be present, but it appears to be at least reduced as compared to the simulated prediction for independent sites. The increased frequency of longer-lived doubly bound events in the data suggests that unbinding of the second ligand is slowed, which in the tested models results from a reduction in the unbinding rate and an increased probability for the CNBD to be in the exchanged conformation from which unbinding is infrequent (disallowed in the model). We also could be underestimating this effect given that the longest-lived events may be truncated by bleaching (Supplementary Fig. 14).

Similarly, we compared the latency to binding of the second ligand (2^nd^ latency) after binding the first ligand in our experimental observations with predictions from simulations of independent sites. The distribution of 2^nd^ latencies was shifted to shorter durations in the experimental fcGMP bound series as compared to simulations of independent sites, again suggesting that the binding rate for the second ligand is slightly faster than that of the first (Supplementary Fig. 13). The ratio for the mean 2^nd^ latency of fcGMP binding data and independent simulations suggests that the second ligand binds 1.3-fold faster than predicted for independent sites, similar to predictions in the tested models. Taken together, the observed durations in distinct ligation states and the models both suggest that binding of the second ligand is similar to the first, but that some potential positive cooperativity may confer a few fold increase in the binding equilibria of the second ligand relative to the first.

## Discussion

Although multicolor SM fluorescence has previously been used to resolve binding dynamics for soluble proteins or nucleotides^50^, similar measures for small molecule binding to membrane proteins in native lipids are few. For oligomeric proteins, the ability to resolve dynamics in distinct ligation states (i.e. singly- vs doubly-bound) is crucial for exploring the sequence of binding events and their cooperative interactions. Here, we introduce a combined approach using cell-derived nanovesicles, microfluidics and mmTIRF colocalization SM fluorescence spectroscopy to study single-receptor ligand binding in native lipids. Our approach has several advantages for SM measurements in membrane proteins: 1) Proteins are never removed from their cellular lipid environment. 2) Vesicles can contain the protein in both inward and outward facing orientations, providing access to either extracellular or intracellular receptor sites. 3) Microfluidic liquid handling enables within-experiment solution exchange to readily identify molecules exhibiting specific binding amidst background signals from nonspecific adsorption that challenge typical SM colocalization experiments at the dye concentrations used. 4) Expression and on-chip purification of full-length proteins for SM experiments requires only standard transfection of cultured cells as opposed to potentially costly and time-consuming searches for appropriate purification conditions in synthetic environments and stabilizing mutations. 5) Immobilization of proteins simplifies the observation of individual molecules over longer time periods to resolve dynamics. As an exemplar system, we applied this approach to study the first two cyclic nucleotide binding steps to full-length CNG TAX-4 channels. Our results suggest that binding is followed by a conformational change of the bound complex similar to that observed in isolated CNBDs from HCN2 channels. Furthermore, our observations provide an unprecedented view of the dynamics of the first two binding events which place constraints on the degree of binding cooperativity between the first and second ligand.

The observed binding dynamics at individual CNBDs in TAX-4 are very similar to observations of fcAMP binding dynamics at isolated CNBDs from HCN2 channels^49^. Coupling these dynamics with structures of the apo and holo CNBD, we previously postulated a mechanism whereby cyclic nucleotide binding is followed by a conformational change of the CNBD involving capping of the binding pocket by the C helix, thereby hindering unbinding prior to uncapping^49^ (Fig. 6a). Such a movement of the C helix is inferred from numerous structures of homologous CNBDs and also observed in functional CNBDs with Förster resonance energy transfer (FRET)^51^ and double electron-electron resonance (DEER)^52^. This mechanism explains the lower frequency of binding/unbinding predicted to occur following the conformational change (e.g. compare models M1.E and M1.F in Supplementary Fig. 9). Given the similarity between our prior observations of purified CNBDs and our observations in this study of full-length TAX-4 channels, it is likely that similar intrinsic fluctuations of isolated CNBDs occur in the channel complex. A recent study arrives at a similar conclusion that a conformational change follows binding in full-length HCN1 and HCN2 channels ^53^.

Cryo-EM structures of TAX-4 have recently been resolved in both unliganded and fully cGMP-bound states^34–36^. These structural snapshots are largely consistent with the above mechanism involving movement of the C helix towards the occupied binding pocket. In the full-length channel structures, binding is also associated with movement of the CNBDs towards the lipid membrane and rotation of the CNBDs about the pore axis resulting in an expansion of the pore lining S6 helices consistent with opening of the channel pore in response to ligand. Similar movements were also observed for CNG channels with fluorescence^39^, DEER^40^, and high-speed atomic force microscopy^41^.

The most likely models explored here predict that the probability of the CNBDs to adopt their alternate conformation increases with successive binding of each of the first two ligands, suggesting that the conformational change is part of the ligand activation pathway. However, it is important to keep in mind that our observations don’t include channel current, which challenges direct comparison with previous models of CNG channel pore gating. Nonetheless, the dynamics of the conformational change at the CNBDs is too slow to account for single channel observations of pore gating^54^, and thus we hypothesize that it reflects an earlier step in the activation process preceding opening of the pore gate. As such, changes in CNBD conformation would place the channel in a preactivated state from which bursts of channel openings could occur. CNBDs in mixtures of conformations could also underlie observed subconductances in channels covalently locked into distinct ligand-bound states^55^. We hypothesize that this conformational change involves capping of the CNBD by the C helix along with movement towards the membrane of the CNBD and C-linker connecting the CNBD to the pore lining S6 helix as observed in structural and functional studies^34–36, 39–41^.

Although our observations of binding dynamics strongly support a conformational change in ligand-bound CNBDs, it is unclear whether this is a global change involving all subunits or independent exchange within individual CNBDs. Models describing both of these processes are difficult to distinguish with these data alone (**Fig. 6b,c**), and likely will require observations of the third and fourth binding steps to resolve. Nonetheless, the dynamics do suggest that binding/unbinding is unfavorable following the conformational change. In the case of a global conformational change, this would also inhibit association at unoccupied sites. Of interest is a model that extends a simple MWC mechanism for ligand binding and pore opening^56^ to postulate that the tetrameric channel behaves as a dimer of dimers^57^. However, given that we only observed the first two binding events, it is difficult to distinguish between channels functioning as tetramers or coupled dimers.

Combined simultaneous observations of fcGMP binding and channel current in macroscopic ensembles of CNGA2 channels previously led to a model with a structure similar to that shown in Fig. 6b, but where the transitions we associate with CNBD conformational exchange were associated with pore gating^22^. This model predicts strong negative cooperativity exclusively during binding of the second ligand and similarly strong positive cooperativity upon binding the third ligand, with a reduction/increase in binding equilibrium of over three orders of magnitude for the second/third binding steps, respectively. Our results rule out such strong cooperativity for the second binding step in TAX-4, rather indicating that binding of the second ligand may involve at most relatively weaker positive cooperativity. Either CNGA2 channels behave differently than TAX-4, or models with more modest cooperative effects^58^ are more likely. We hypothesize that occlusion of individual binding events in distinct ligation states by ensemble averaging challenged unique identification of cooperative effects. Ultimately, additional experiments are needed to resolve the dynamics of the third and fourth binding steps, possibly with FRET between fcGMP and an acceptor label on the channel to retain SM resolution at higher fluorophore concentrations^59^. A recent study using similar single-molecule methods shows that cAMP binding is non-cooperative in HCN1 and HCN2 channels^53^. However, whereas CNG channels are directly gated by cGMP, HCN channels are gated primarily by voltage and modulated by cAMP. Thus, although our observations for TAX-4 are qualitatively similar to that for HCN1 and HCN2, positive cooperativity for the second binding step may reflect ligand-induced pore opening in TAX-4, although this remains to be determined.

This study provides unprecedented observations of the dynamics of early binding events and a conformational change that precedes channel opening. Our observations complement static structural snapshots of fully unliganded and fully bound conformations by reporting on the dynamics of partially liganded conformations that lie on the reaction pathway between endpoints designated by recent cryo-EM structures. These data also exemplify how binding dynamics can be used to observe conformational exchanges that may otherwise be difficult to measure, either due to their transient nature, small structural motions, or because it is unclear where to place probes for more targeted measurements. Finally, our combined SM approach has broad application to any membrane protein where protein and/or ligand are amenable to fluorescent labeling.

## Methods

### Constructs

TAX-4 was a gift from Drs. Jonathan Pierce and Iku Mori. EGFP was appended to the N-terminus of TAX-4 (GFP-TAX-4) in the pUNIV vector as follows: A previous construct with an N-terminal EGFP in pUNIV containing an ApaI restriction site between EGFP and the gene and an MluI restriction site following the gene was used as a starting template. The previous gene was excised between ApaI and MluI sites, and TAX-4 was inserted using the same sites. This resulted in a two-residue linker (G-P) between EGFP and TAX-4. The entire gene was sequenced for verification.

The full-length rat GABA_A_ receptor α1, β2 and γ2L subunits in the pUNIV vector were a gift from Dr. Cynthia Czajkowski. mScarlet (Addgene #99280) was inserted in the M3-M4 loop of the α1 subunit between residues V372 and K373 using an in-frame non-native Asc1 restriction site (GGGCGCGCC) introduced through site-directed mutagenesis as previously described^60^. This results in the insertion of an additional three residues (G-R-A) on each end of mScarlet. EGFP was similarly inserted in the N terminal region of the β2 subunit between residues N4 and D5, again using an in-frame non-native Asc1 restriction site as described for mScarlet.

### Preparation of cell-derived nanovesicles

HEK-293T cells were cultured at 37 °C and 5% CO_2_ (Eppendorf). Cells were plated in 60 mm dishes and transfected with 1 ug of GFP-TAX4 and 3 ug of PEI-MAX per dish. After 24 hours, cells from four dishes were combined and vesicles were prepared as previously described^9^. Briefly, the cells were subject to nitrogen cavitation at 600 psi. for 20 minutes while suspended in 3 mL of hypotonic protease inhibitor solution (in mM: 10 Tris-HCl, 10 NaCl, 1.5 MgCl2, 0.2 CaCl2, pH 7.4). One Pierce protease inhibitor tablet (ThermoScientific) was added per 10 ml of buffer. To separate plasma membrane vesicles from organelle membrane vesicles, the lysate was dispensed onto a gradient containing 60, 30, 20, and 10% solutions of OptiPrep, followed by ultracentrifugation at 112,000 x g for 90 min at 4 °C. Following centrifugation, the sample was fractionated into nine 1-1.5 mL fractions using a peristaltic pump, where the highest density fraction was collected first. OptiPrep was removed from fractions containing plasma membrane nanovesicles via centrifugation at 100,000 x g for 1 hour at 4 °C using a fixed angle rotor. The resulting pellet was resuspended in 250 μL of buffer for immobilization as described below.

### Imaging fcGMP binding at TAX-4 channels in cell-derived nanovesicles

Prior to immobilization, coverslip glass was UV-cleaned (Jelight) and passivated with a PEG monolayer sparsely doped with PEG-biotin (Laysan Bio) as previously described^44, 61^. A 50 μL microfluidic chamber (Grace Biolabs) was adhered to the passivated coverslip to allow for solution flow during imaging. The chamber was serially incubated in 10 mg/ml bovine serum albumin (BSA), 50 ug/ml streptavidin, and 1 ug/mL biotinylated GFP-nanobody (ChromoTek). All solutions were made in a buffer that consisted of phosphate buffered saline (PBS; pH 7.4) supplemented with 1 mM CaCl_2_ and 1 mM MgCl_2_. Following each 10-15 minute incubation step, the chamber was washed with 4 ml of buffer to remove any non-immobilized components. Finally, the chamber was incubated with a preparation of nanovesicles from cells expressing GFP-TAX-4 (i.e. protein of interest fused with GFP) for 10-15 minutes followed by washing with 4 ml of buffer. This resulted in sparse immobilization of vesicles containing GFP-TAX-4. The chamber was then connected to a microfluidic pump and switch (Elveflow) for perfusion and exchange of solutions containing various concentrations of 8-(2-[DY-547]-aminoethylthio) guanosine-3’,5’-cyclic monophosphate (fcGMP; BioLog) and cGMP in buffer. Solutions were continuously perfused at a constant flow rate of approximately one chamber volume per minute during imaging on an inverted mmTIRF microscope (Mad City Labs) under either 488 or 532 nm laser excitation (Coherent OBIS). Laser power at the sample was 20 W/cm^2^ (488 nm) and 40 W/cm^2^ (532 nm). Fluorescence emission from immobilized vesicles in response to mmTIRF excitation was recorded simultaneously for an ∼100×100 μm field of view on a 512×512 EMCCD camera (Andor iXon) at a frame rate of 20 Hz (50 ms per frame). For each field of view, we initially determined the location of all immobilized vesicles by bleaching the EGFP emission from 500-550 nm under 488 nm excitation. Thereafter, binding of fcGMP in solution was monitored by recording emission from fcGMP within 560-950 nm under 532 nm excitation.

### Single-molecule fcGMP binding image analysis

Time-averaged fluorescence for EGFP and fcGMP from the same field of view were overlaid to identify diffraction-limited spots containing GFP-TAX-4 that colocalized with fcGMP binding. For EGFP this average included only the first 20-50 frames prior to significant bleaching of the EGFP signal. The entire time series was averaged for fcGMP as unbleached fcGMP from the bath can continuously diffuse into the vesicle layer where binding is detected. Mechanical drift of the stage parallel to the imaging plane was subpixel during imaging even for tens of minutes or longer due to the high stability of the Mad City Labs mmTIRF stage. A small offset of up to a few pixels was sometimes observed between EGFP and fcGMP image sets due to mechanical perturbation from manual swapping of filters on the optical table. This offset was corrected for by registering images of the time-averaged fluorescence for EGFP and fcGMP recordings in MATLAB using an affine transform. Drift perpendicular to the image plane was continuously autocorrected during imaging by a nano-positioning stage that adjusted the distance between the sample chamber and microscope objective to maintain the position of the mmTIRF excitation laser on a quadrant position detector downstream of the exit micromirror (Mad City Labs). For colocalized spots with both EGFP and fcGMP fluorescence, the average intensity within a five-pixel diameter circle centered on the spot was projected across each frame to obtain the fluorescence time series at that spot. Bleach steps for EGFP at each and every colocalized spot were manually evaluated to estimate the number of GFP-TAX-4 subunits in each spot and spots with more than four EGFP bleach steps consistent with multiple channels were excluded from analysis. The number of molecules at each concentration was: 10 nM: 112, 30 nM: 325, 60 nM: 63, 100 nM: 65 and 200 nM: 132. In total, we recorded ∼60 hours of single-molecule binding.

To assess the impact of photobleaching on the observed event durations we examined the lifetimes of noncolocalized fcGMP events that we assume largely reflect adsorption to the surface and subsequent bleaching. If some of these events are truncated by unbinding from the surface rather than bleaching, we will underestimate the actual bleach times. Given that these events were typically noisy, we manually estimated their lifetimes for a handful of randomly selected experiments. The mean bleach time across molecules was 23.4 seconds, an order of magnitude longer than the time constant for the longest duration bound component (Supplementary Fig. 14; Supplementary Table 1). This suggests that most binding events were terminated by unbinding rather than bleaching, although the lifetimes of the longest-lived bound events are likely to be slightly underestimated due to bleaching.

### Idealization of time series for number of fcGMP bound

To extract time series for the number of bound fcGMP we first denoised fcGMP fluorescence time series using DISC^62^. A complication in interpreting these time series is that the fluorescence intensity for individual events was somewhat variable, challenging assignment of the number of bound ligands at each time point based solely on intensity. The distribution of individual event intensities is close to normally distributed with a slight skew toward higher intensities (Supplementary Fig. 7), suggesting that this variation reflects a random process rather than distinct numbers of bound ligands or discrete subpopulations of receptors. Such variation is typically observed in other colocalization fluorescence experiments. The source of this heterogeneity is uncertain but is likely caused by shifts of the molecule in the exponentially decaying excitation field or dye photodynamics^63, 64^. To address overfitting due to fluctuation of individual event intensities, neighboring piecewise constant segments identified by DISC were recursively merged if the difference between their mean fluorescence was less than the weighted sum of the standard deviations of their fluorescence, where the weight for each segment was the ratio of the number of data points in the segment to the total number of data points in both segments. Single frame segments with intensities intermediate to their surrounding segments were visually identified as typically reflecting noise rather than discrete transition intermediates, and thus were merged into the neighboring segment with the most similar mean intensity. The denoised fluorescence time series was then constructed from the mean fluorescence within each discrete segment. The lowest intensity level identified by DISC was assumed to reflect the unliganded baseline, as even at the highest concentration tested periods of low fluorescence consistent with background were observed for all molecules. Any denoised segment containing any of these baseline data points was assigned a bound count of zero. For each contiguous block of remaining segments, the number of bound fcGMP was incremented or decremented based on whether the denoised intensity increased or decreased. Binding/unbinding events whose denoised intensity change was more than double that of neighboring unbinding or binding events, respectively, were considered to involve binding or unbinding of two molecules of fcGMP within a single frame. Similarly, contiguous triplets of events that returned to within 100 au of their initial intensity were considered to include a single event for association or dissociation of two fcGMP within a single frame. In a few cases this procedure resulted in a bound count of less than one for a segment we previously identified as having a nonzero number of bound ligands, suggesting that we misidentified the number of fcGMP associated with a prior individual event. To correct for this, we recursively examined the chain of monotonic unbinding events preceding each segment with an erroneous bound count of zero. If this chain contained a segment with only a single frame, the two surrounding unbinding events were merged into a single unbinding event, otherwise the number of fcGMP associating during the binding event preceding this chain was incremented by one. This resulted in an idealized series for the number of bound fcGMP from 0-4 at each time point.

To assess the ability of the idealization procedure to resolve individual events, we applied it to simulated binding data at 30 or 200 nM ligand concentration whose lifetimes and noise were drawn from the experimental fluorescence observations (see below and Supplementary Fig. 7). Comparison of the known and idealized event records for simulated binding data indicates that the idealization procedure is incredibly accurate at identifying singly and doubly liganded events lasting two or more frames (>= 100 ms) but misses 36% brief single frame dwells in doubly liganded states (Supplementary Fig. 8). This is due to the additive noise from multiple fluorescent ligands coupled with a dwell time on the order of our sample duration and is much less of an issue for single frame events with only one occupied site (10% missed). Thus, brief single-frame intensity fluctuations following binding of one or more ligands that were identified as noise during idealization (e.g. see couple of spikes around 15 seconds in Fig. 4b) could include missed brief events at higher ligation states. The algorithms overall accuracy, precision and recall was computed allowing for misidentification of the exact timing of individual events by up to four frames (accuracy = 0.83, precision = 0.95, recall = 0.87, F1-score = 0.91).

### Simulated fcGMP binding time series

To simulate binding data resembling our experimental recordings, we used the idealized bound time series (see above) to extract baseline Gaussian noise and distributions of bound and unbound dwell times as well as isolated binding event intensities and their Gaussian fluctuations. To address periods with multiple bound ligands, we estimated individual site bound dwell times by randomly assigning each unbinding event to one of the bound ligands. Unbound dwell times for individual sites were assumed to be on average four times longer than were observed at tetrameric channels with four sites.

We first simulated single-site bound time series by randomly drawing from the single-site dwell time and event amplitude distributions described above, followed by the addition of Gaussian noise to individual event segments. Gaussian noise for individual events was assigned based on the observed linear correlation between event intensity and the standard deviation of within-event fluctuations for events with duration longer than 10 frames. No noise was added to the baseline at this point. To simulate tetramers comprised of independent CNBDs, we added four single-site time series together, and finally added Gaussian noise to the remaining baseline points based on the average standard deviation of baseline fluctuations in our data. This procedure sums the noise from each bound ligand without quadrupling the baseline noise. In total, we simulated ∼60 hours of binding data at multiple fcGMP concentrations to obtain a simulated dataset similar in size to our experimental dataset.

### HMM Analysis

All models were optimized in QuB^47, 48^ to maximize their likelihood for ∼60 hours of idealized binding events across all molecules and concentrations. For single-site models (Supplementary Fig. 9) we restricted our analysis to isolated binding events by removing periods with two or more bound ligands, thereby splitting those time series into segments comprised only of singly bound events. For models of the first and second binding steps (Supplementary Fig. 10), we similarly restricted our analysis to periods with up to two occupied sites by removing bound periods with more than two bound ligands. Dead time in all cases was one frame. Optimized rate constants and their estimated uncertainty are given in Supplementary Tables 2 and 3. The models were ranked according to their relative Bayesian Information Criterion (*Δ*BIC = BIC – BIC_best model_) scores (smaller is better) (Supplementary Figs. 9-10; Supplementary Tables 2-3).

## Data availability

The time series data that support the findings of this study are available as supplementary material. Additional data such as original image series are available from the authors upon request.

## Code availability

Data analysis was performed using custom-written scripts in MATLAB available as supplementary material and at https://github.com/marcel-goldschen-ohm/single-molecule-imaging-toolbox.

## Acknowledgements

We thank Drs. Jonathan Pierce and Iku Mori for gifting cDNA for wild-type TAX-4. We thank Drs. Richard Aldrich and Eric Senning for helpful discussion and shared use of equipment and Dr. Eric Senning for gifting the TRPV1-GFP construct.

## Author Contributions

V.R.P. performed molecular biology including generating cDNA for constructs, carried out the experiment, analyzed the data and contributed to writing of the manuscript. A.M.S. and D.Q. contributed to data analysis and writing of the manuscript. S.G. performed molecular biology including generating cDNA for constructs as well as all cell culture for experiments. D.J.S. performed preliminary vesicle experiments. M.P.G. conceived, designed and oversaw the project and contributed to data analysis and writing of the manuscript.

## Supplementary Material

**Supplementary Fig. 1.**
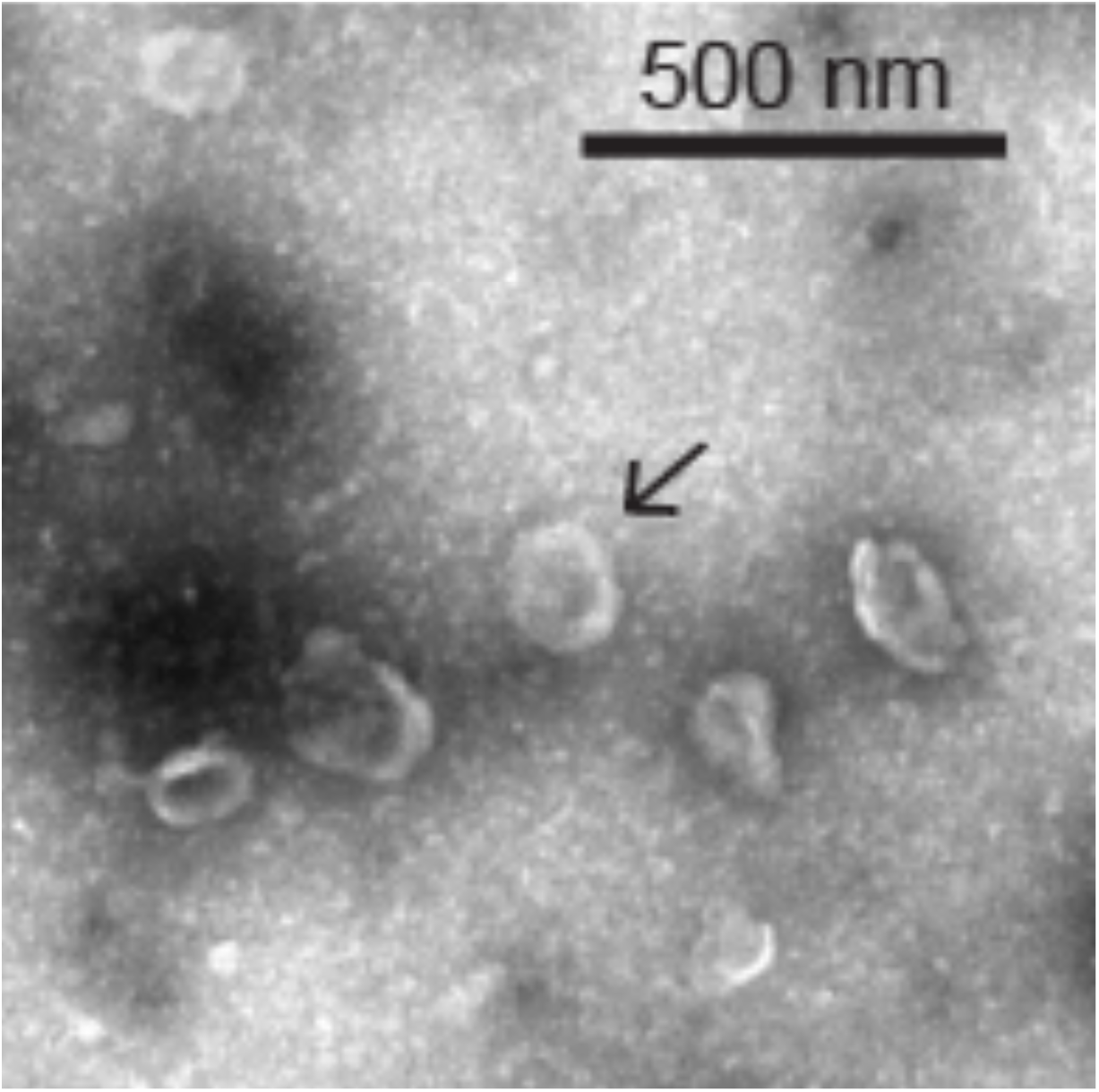
Transmission electron microscopy (TEM) images of cell-derived nanovesicles. Vesicle diameters in TEM images of a standard vesicle preparation (see Methods) ranged from approximately 50–1000 nm in diameter similar to previous reports based on dynamic light scattering^1^. Arrow indicates a single vesicle with a diameter of ∼150 nm.

**Supplementary Fig. 2.**
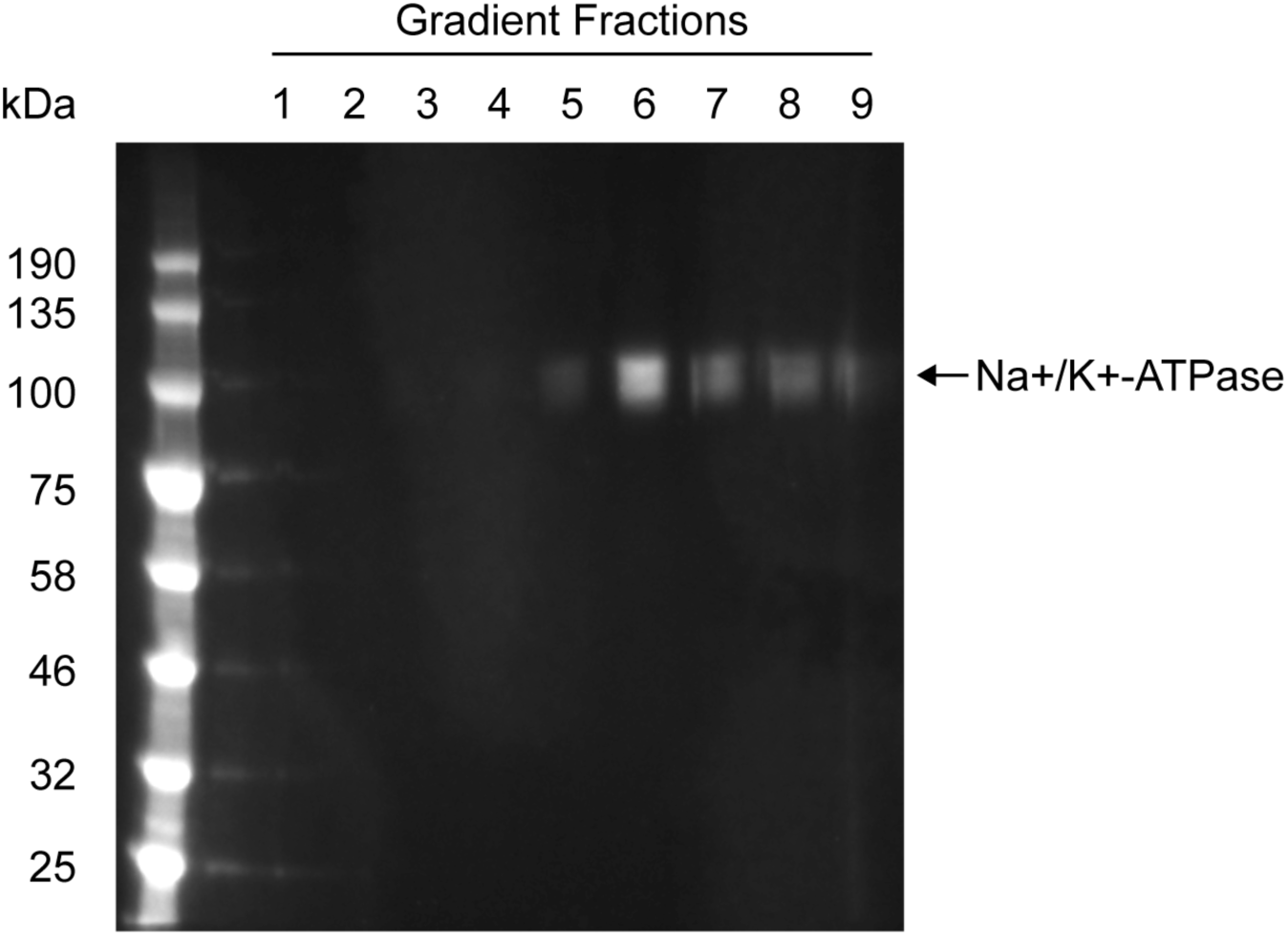
Western blot of plasma membrane fraction labeling. The left-most lane corresponds to molecular weight markers used as a standard. The same amount of nanovesicle lysate from fractions 1-8 following density gradient ultracentrifugation was loaded into the corresponding wells. Fractions containing nanovesicles derived from plasma membrane were detecting using fluorescent antibodies specific to Na+/K+-ATPase, which were enriched in plasma membrane fractions 5-8. These data confirm an earlier report^2^.

**Supplementary Fig. 3.**
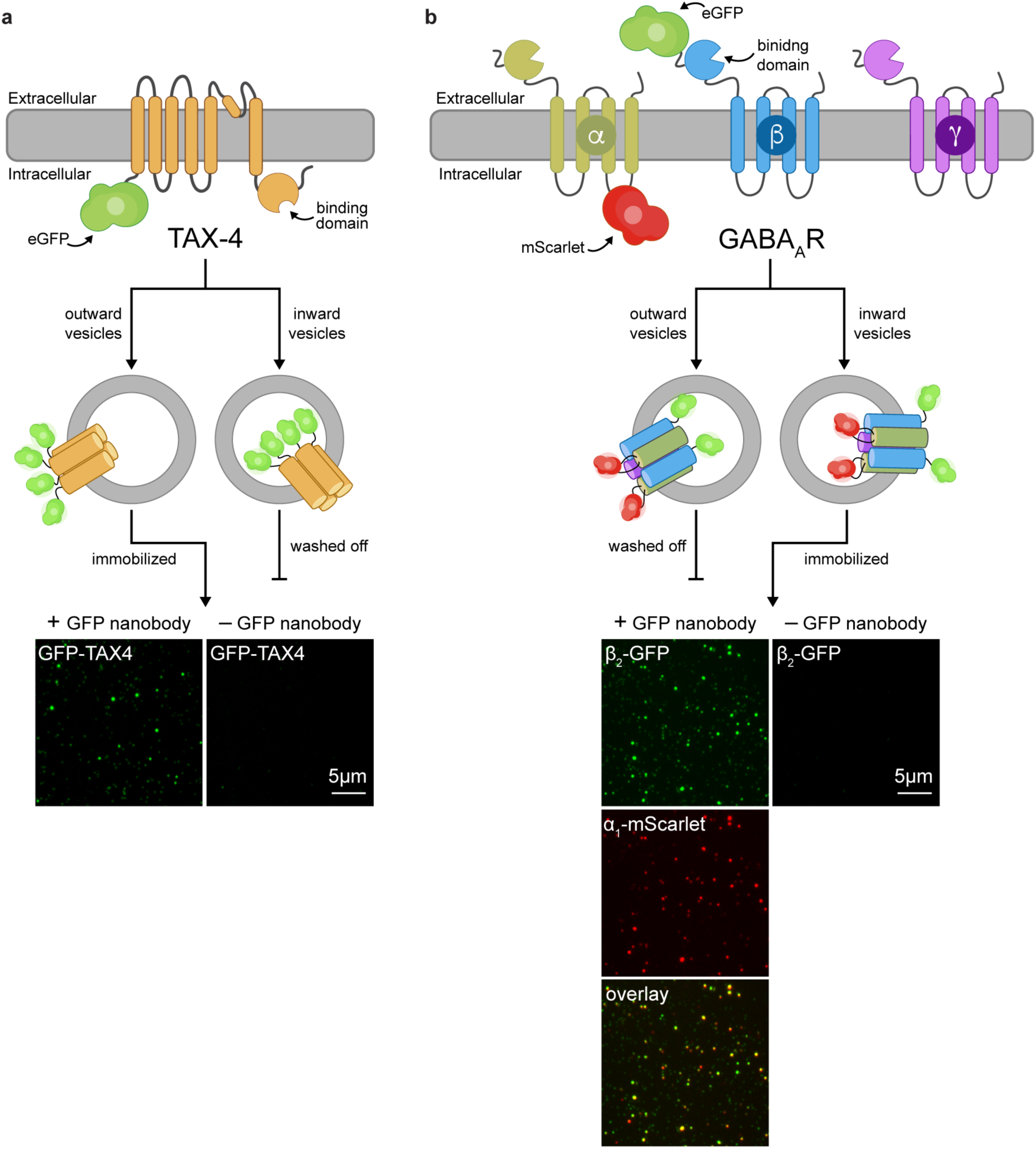
Specific immobilization of nanovesicles containing a membrane protein of interest. **a** Cartoon illustrating topology of a GFP-TAX-4 subunit with intracellular cyclic nucleotide binding domain and vesicles derived from cells expressing GFP-TAX-4 tetrameric channels. Below are mmTIRF images of GFP fluorescence (green) showing specific immobilization of GFP-TAX-4 containing vesicles at GFP-nanobodies deposited on the imaging surface (left) and a lack of appreciable nonspecific surface adsorption of vesicles in the absence of GFP-nanobodies (right). In both cases chambers were incubated in GFP-TAX-4 containing vesicles and then rinsed with buffer before imaging. **b** Cartoon illustrating topology of GABA_A_ receptor *α*_1_, *β*_2_ and *γ*_2L_ subunits and vesicles derived from cells expressing heteropentameric channels. For visualization and immobilization mScarlet was inserted in the intracellular M3-M4 linker of *α*_1_ (*α*_1_-mScarlet) and EGFP was inserted near the N-terminus of *β*_2_ (*β*_2_-GFP) on the same side of the membrane as the extracellular agonist and benzodiazepine binding domains. Below are mmTIRF images of GFP (green) and mScarlet (red) fluorescence showing colocalization (yellow) of *β*_2_-GFP and *α*_1_-mScarlet subunits at vesicles immobilized with GFP-nanobodies and a lack of appreciable nonspecific surface adsorption of vesicles in the absence of GFP-nanobodies.

**Supplementary Fig. 4.**
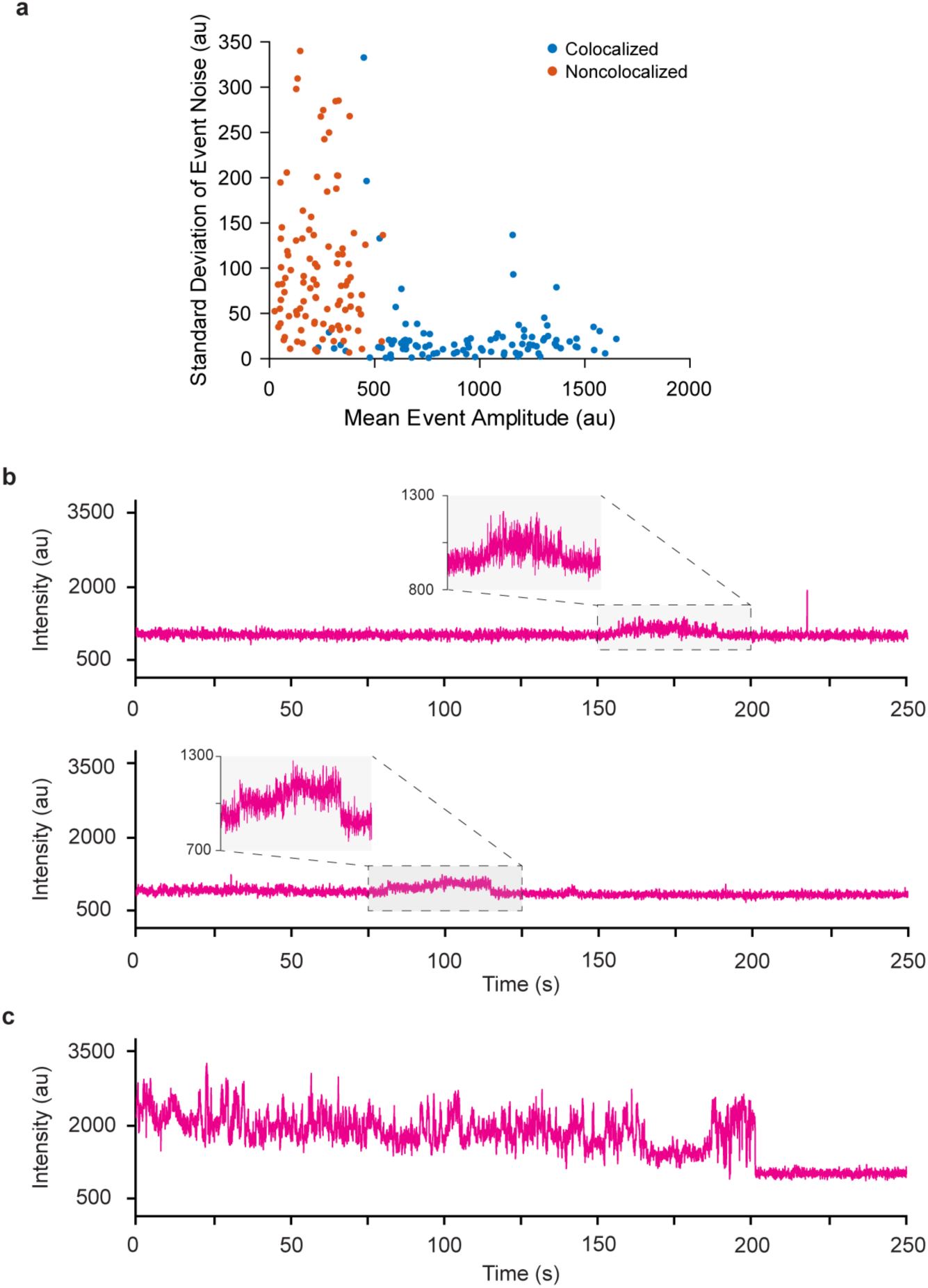
Distinct properties between colocalized and non-colocalized fcGMP binding events. **a** A comparison of the mean and standard deviation of the fluorescence signal from individual fcGMP binding events at spots that either do or do not colocalize with GFP-TAX-4 in a single field of view. The non-colocalized binding events are likely due to fluorescent contaminants in the PEG layer or non-specific adsorption of fcGMP to imperfections in the surface passivation. **b** Representative traces for low-intensity non-colocalized fcGMP signals, which make up a majority of the nonspecific signal. **c** Representative trace for high-intensity non-colocalized fcGMP signal likely due to non-specific adsorption to glass imaging surface. These high intensity traces make up a minority of observed non-colocalized signals.

**Supplementary Fig. 5.**
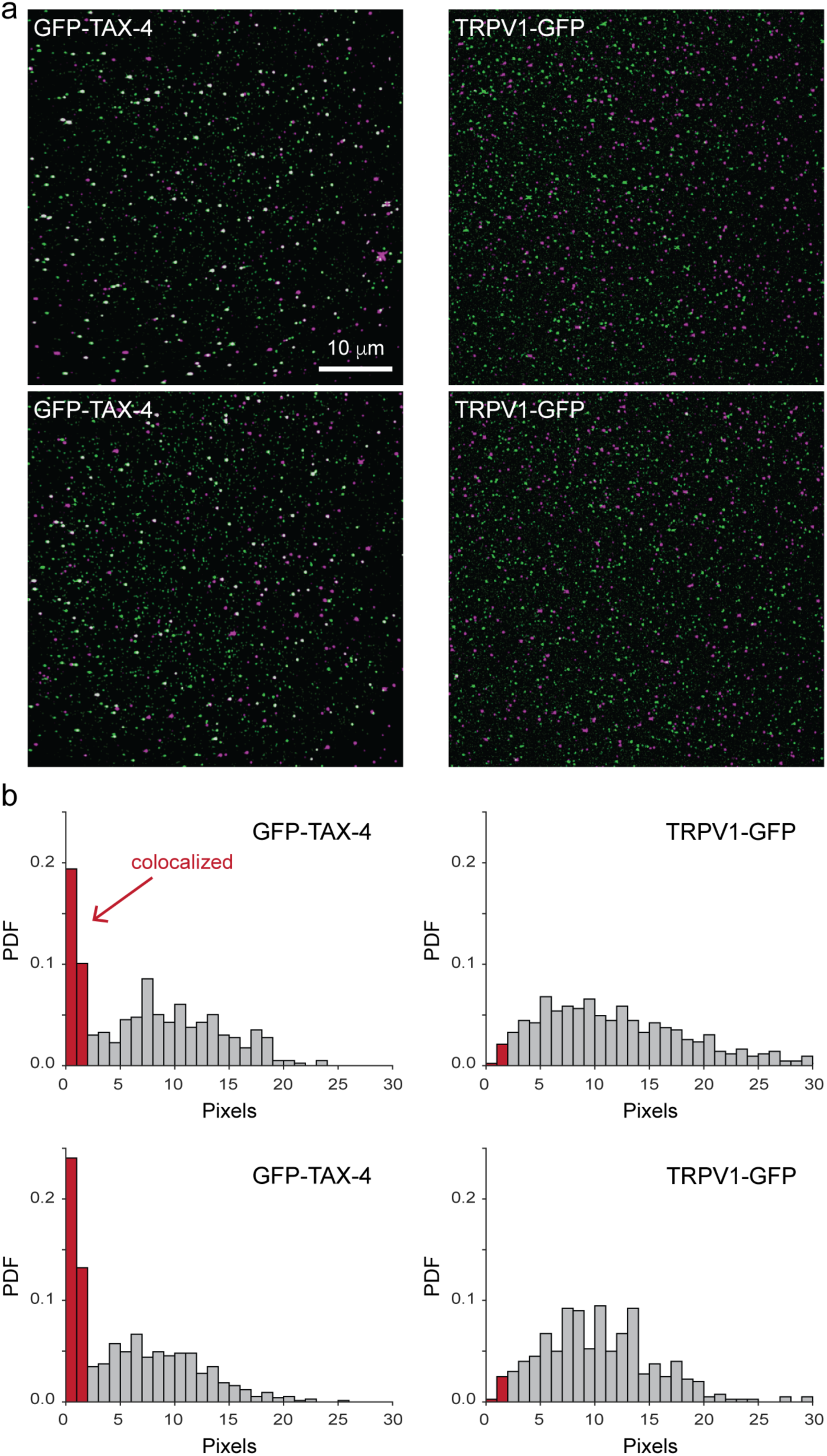
Summary of GFP and fcGMP colocalization for vesicles with GFP-TAX-4 and controls for vesicles with TRPV1-GFP channels. **a** Time averaged fluorescence for GFP (green) from nanovesicles containing either GFP-TAX-4 (left) or TRPV1 channels fused with an intracellular GFP (right) overlaid with 10 nM fcGMP (magenta) from the same field of view (two separate imaging locations shown for each construct). Colocalized GFP and fcGMP signals appear white. Note the relative absence of colocalization for TRPV1 channels as compared to TAX-4 channels. Also note that some of the fainter green spots are subthreshold as only the brighter more distinct spots were observed in the presence but not absence of GFP. **b** Histograms of the nearest neighbor distance from each identified GFP location to the closest identified fcGMP location. Two separate examples are shown for both GFP-TAX-4 and TRPV1-GFP vesicle preparations. The percentage (mean ± standard deviation) of GFP spots that colocalized with fcGMP spots (i.e. were within 3 pixels) across the dataset were for GFP-TAX-4 (at each fcGMP concentration): 33 ± 9% (10 nM), 32 ± 9% (30 nM), 31 ± 7% (60 nM), 33% (100 nM), and 39 ± 3% (200 nM). For TRPV1-GFP at 10 nM fcGMP a background colocalization of 7 ± 0.5% was observed.

**Supplementary Fig. 6.**
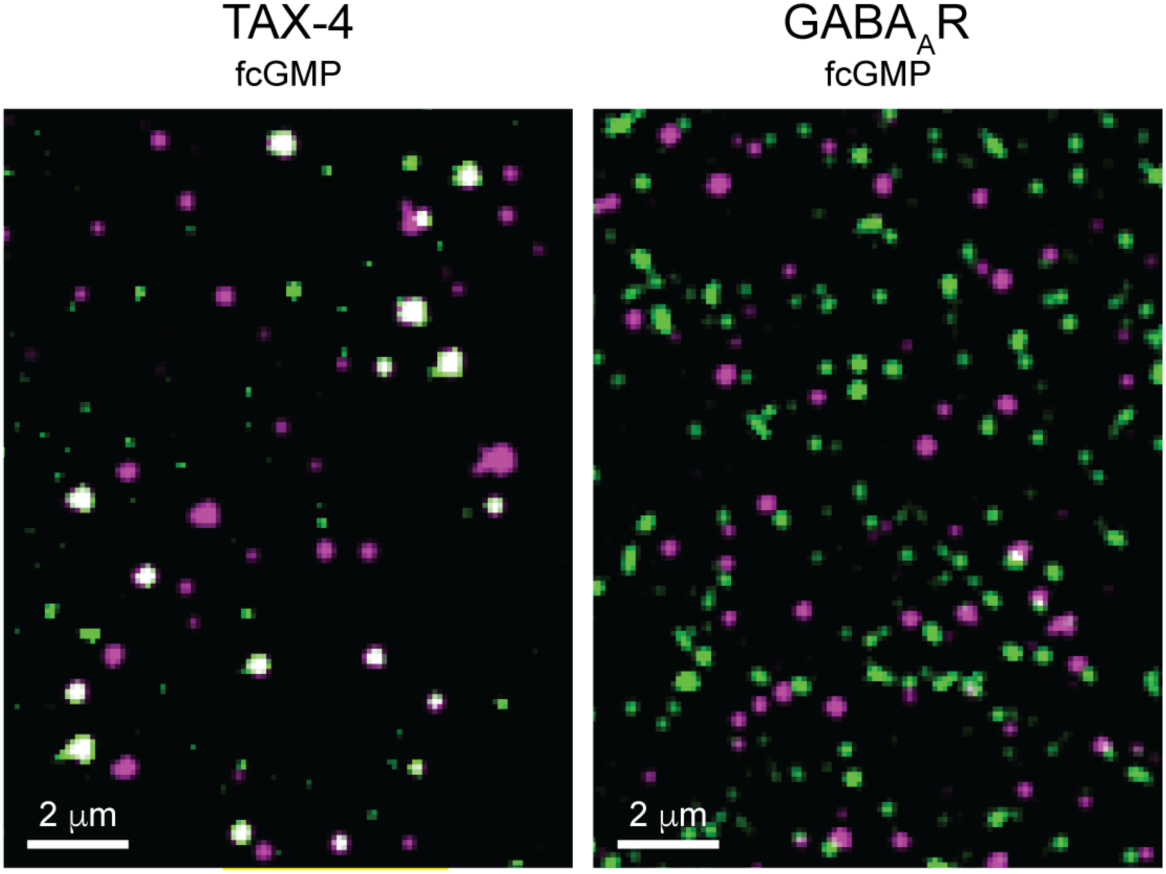
No colocalization between immobilized GABA_A_R containing nanovesicles and fcGMP. Time averaged fluorescence for GFP (green) from nanovesicles containing either GFP-TAX-4 (left) or GABA_A_ receptor *α*_1_, *β*_2_ and *γ*_2L_ subunits with GFP inserted in the N-terminus of the *β*_2_ subunit (*β*_2_-GFP) (right) overlaid with 100 nM fcGMP (magenta) from the same field of view. Colocalized GFP and fcGMP signals appear white. See Supplementary Fig. 3 for a description of the GABA_A_R subunits.

**Supplementary Fig. 7.**
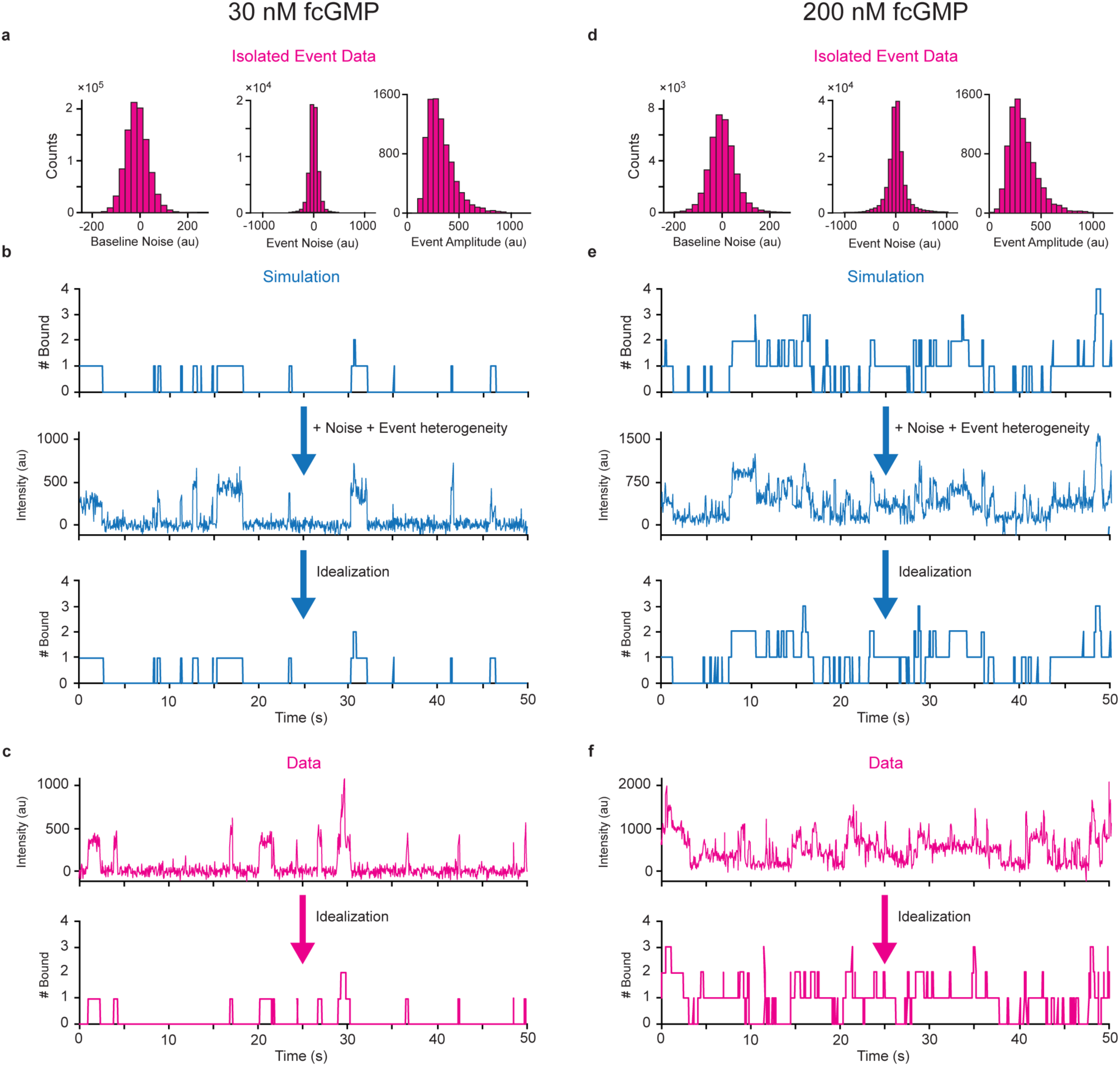
Evaluating the accuracy of idealization of fluorescence time series for ligand binding using simulated data. **a** Distributions of baseline noise, event noise, and event amplitudes for experimental binding data from single, isolated events at 30 nM fcGMP. **b** Example of time series from independent site binding simulations, followed by addition of gaussian noise and event amplitude heterogeneity representing isolated event data (a), followed by the results of our idealization procedure. Comparison of the known simulated time series to the results of our idealization procedure after adding noise drawn from experimental observations provides a metric for testing the accuracy of our idealization procedure (see Methods in main text). **c** Example of binding data at 30 nM fcGMP, followed by the results of our idealization procedure. **d-f** Same as described for **a-c**, but for simulations and data at 200 nM fcGMP.

**Supplementary Fig. 8.**
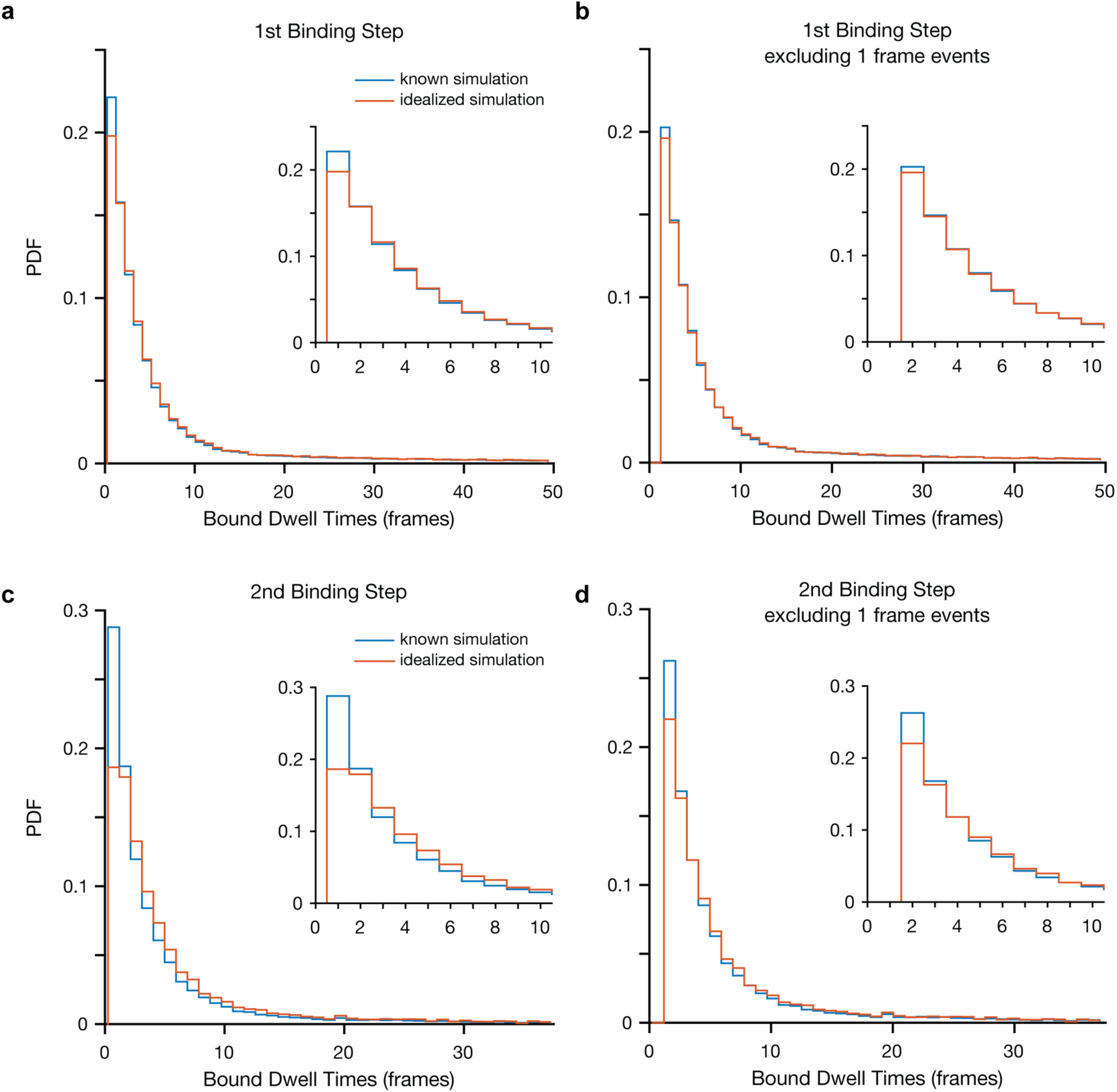
Missed events during idealization of simulated fluorescence time series. **a-b** Simulated bound dwell time distributions of isolated singly bound events either including (a) or excluding (b) single frame events. Note that durations are shown in frames. Comparison of known simulated bound series (blue) to the idealized bound series obtained from idealization after adding experimentally relevant noise and event heterogeneity (red). The primary discrepancy is limited to events lasting only a single frame, which are sometimes missed during the idealization procedure. However, nearly all events lasting two or more frames are reliably detected. **c-d** Same as described for (a-b), but for dwell times in doubly bound states. All simulations were run with a frame rate of 20 Hz (50 ms per frame), equivalent to experimental recordings.

**Supplementary Fig. 9.**
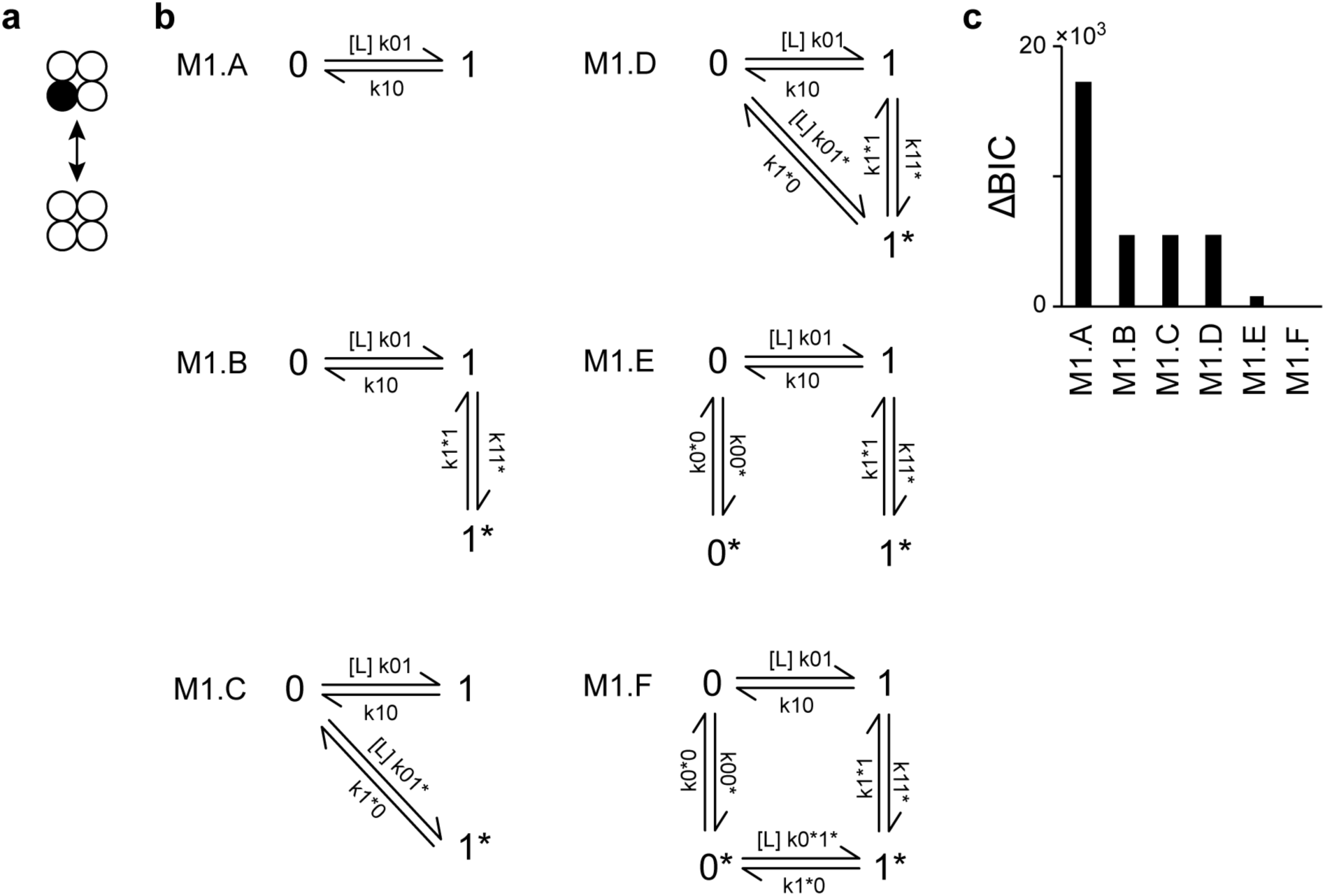
Single site dynamics for the first binding event. **a** Schematic illustrating binding to the first of four sites per channel. CNBDs depicted in unbound (open circles) and bound (filled circle) conformations. **b** Evaluated single site models describing binding and conformational exchange between distinct bound or unbound states. State names indicate the number of bound ligands and an asterix denotes distinct states having the same number of bound sites. Rate constants are in μM^−1^s^−1^ for binding transitions (0 → 1, 0 → 1^∗^, 0^∗^ → 1^∗^) and s^−1^ for all other transitions. [L] indicates ligand concentration for binding steps. Rate constants were optimized in QuB for data segments comprised of isolated binding events (no stacked events) across all molecules and concentrations (see Methods in main text). For models M1.D and M1.F one of the rate constants was constrained to enforce microscopic reversibility in the loop. Rate constants and their estimated errors are given in Supplementary Table 2. **c** BIC scores for the models shown in C relative to the model with the best score (smaller score is better; ΔBIC = BIC – BIC_best model_).

**Supplementary Fig. 10.**
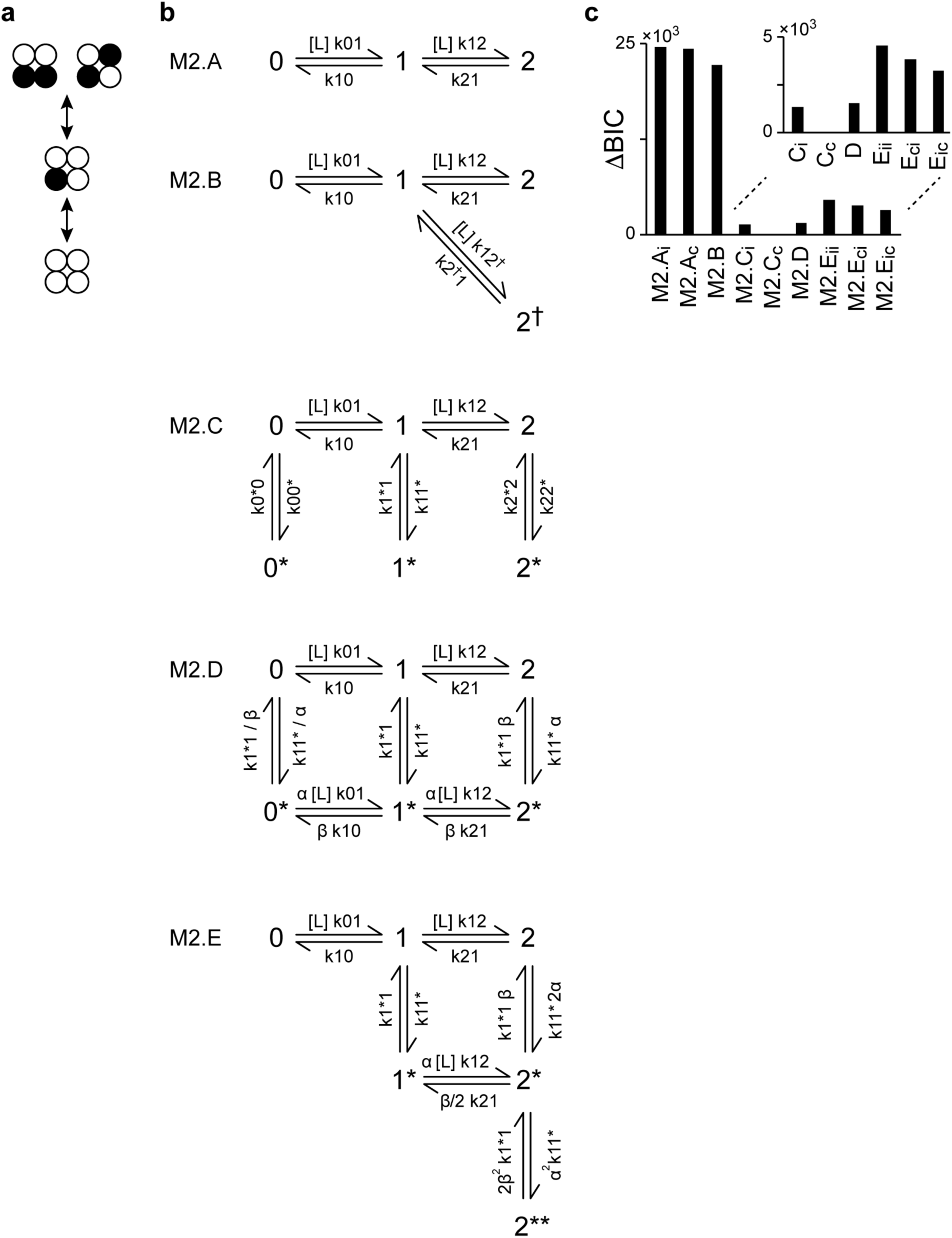
Dynamics for the first two binding events. **a** Schematic illustrating binding to the first two of four sites per channel. CNBDs depicted in unbound (open circles) and bound (filled circle) conformations. **b** Evaluated two site models describing binding and conformational exchange between distinct bound or unbound states. State names indicate the number of bound ligands and an asterisk denotes distinct states having the same number of occupied sites. Rate constants are in μM^−1^s^−1^ for binding transitions (0 → 1, 1 → 2, 1 → 2^†^, 0^∗^ → 1^∗^, 1^∗^ → 2^∗^) and s^−1^ for all other transitions. [L] indicates ligand concentration for binding steps. M2.A is the simplest possible scheme for sequential binding at two sites. M2.B allows for two distinct di-liganded states (e.g. adjacent and diagonally opposed occupied sites as depicted in *a*). M2.C and M2.D extend M2.A with a global conformational change of both sites (between state % and %*). M2.E describes binding followed by a conformational change at each site separately. Note that only model M2.B distinguishes between adjacent and diagonal doubly bound conformations in the channel tetramer. Each model is further subdivided into several models sharing the same schematic but differing in the applied constraints. For example, M2.A_i_ and M2.A_c_ both have the M2.A structure, but A_i_ constrains the two binding steps to be identical and independent, whereas A_c_ allows the dynamics for the two steps to differ (i.e. allows cooperativity between binding sites). Models C_i_ and C_c_ share the same constraints as A_i_ and A_c_. M2.E_ii_ assumes completely independent and identical sites, whereas M2.E_ci_ allows for binding of the 2^nd^ ligand to differ from that of the first (binding cooperativity), and M2.E_ic_ allows the conformational change following binding at either site to depend on the number of bound ligands. See Supplementary Table 3 for a description of all applied constraints. Rate constants were optimized in QuB for the entire data set across all molecules and concentrations excluding events with more than two bound ligands (see Methods in main text). Rate constants and their estimated errors are given in Supplementary Table 3 along with model constraints. **c** BIC scores for each model relative to the model with the best score (smaller score is better; ΔBIC = BIC – BIC_best model_). See description above for model subscripts.

**Supplementary Fig. 11.**
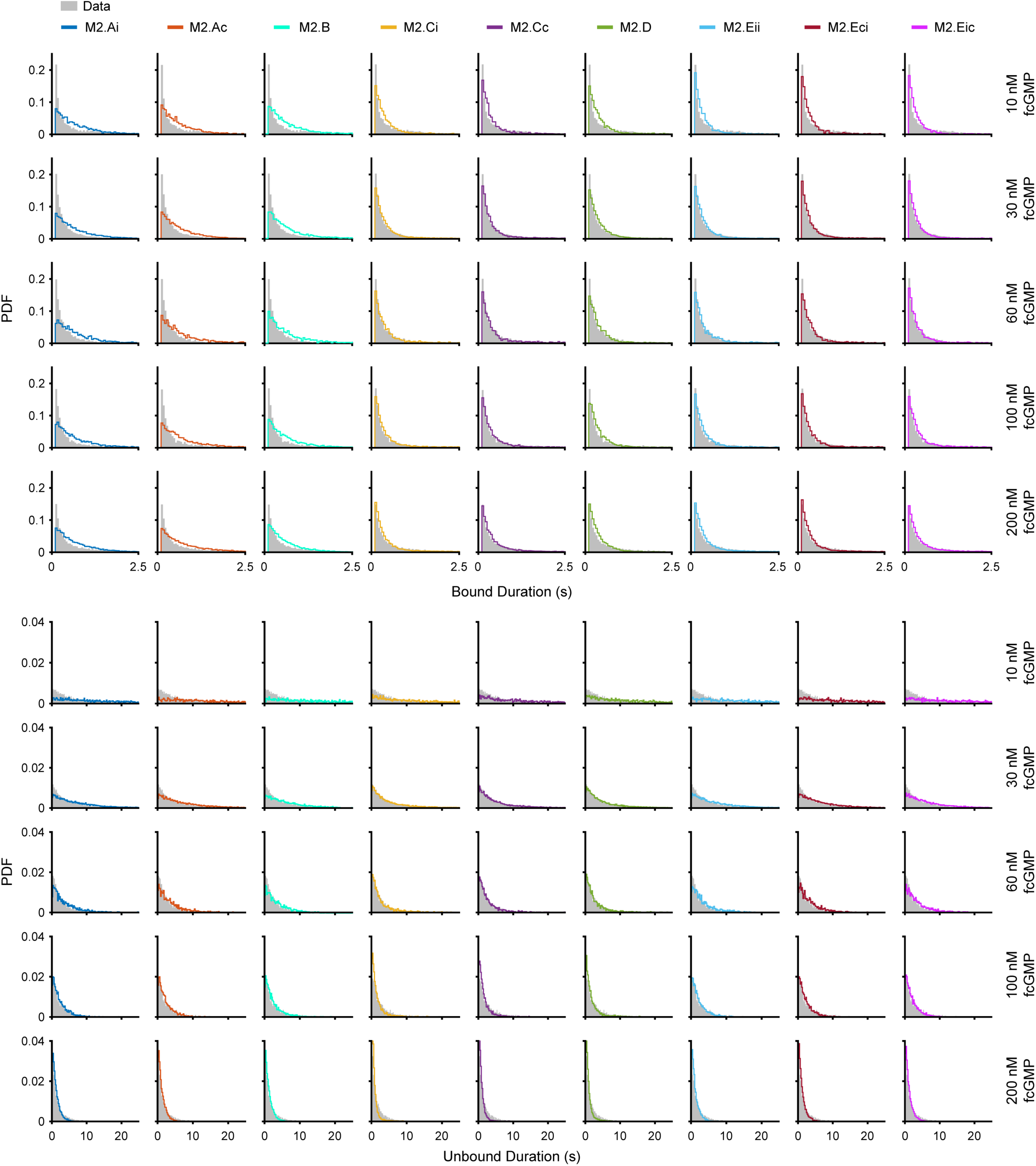
Comparison of observed dwell time distributions with predictions for two-site models. Experimentally observed bound and unbound dwell time distributions (gray) overlaid with predicted distributions from simulated bound time series for each two-site model (colored; see Supplementary Fig. 10). For each ligand concentration, simulations matched the length of data collected.

**Supplementary Fig. 12.**
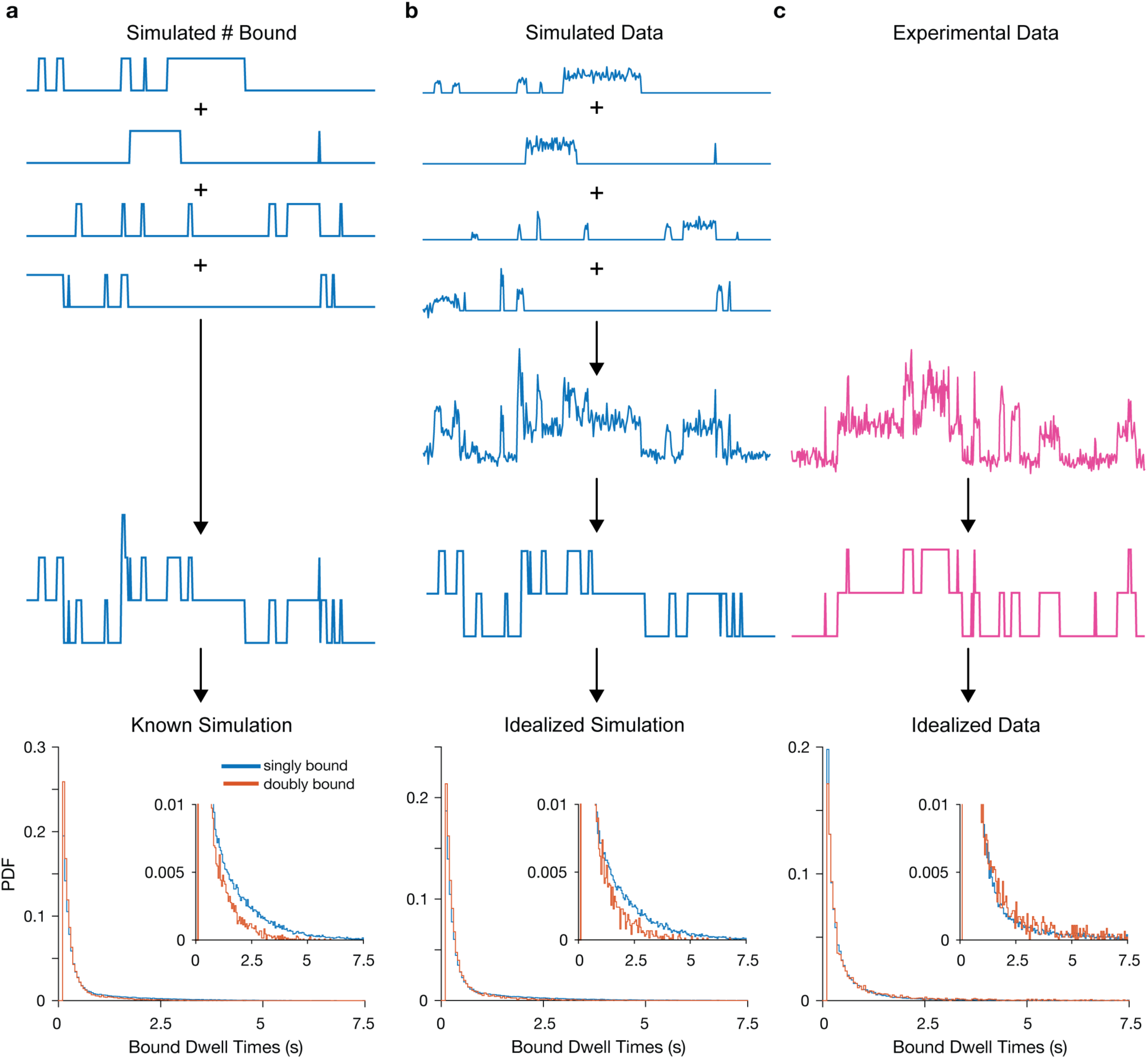
Comparison of experimental data with simulations for independent CNBDs. **a** Four simulated single-site bound time series were added together to generate bound seeries representative of tetramers comprised of independent CNBDs. **b** Gaussian noise and event amplitude heterogeneity were added to simulated bound series to reflect experimental observations, and the noisy simulations were idealized to obtain bound series. **c** Experimental fcGMP binding fluorescence traces idealized to obtain estimated bound series in exactly the same way as for the simulated data in (b). Dwell time distribution in singly bound (blue; one of four CNBDs occupied) and doubly-bound (red; two of four CNBDs occupied) states are shown for both known simulations (a), idealized simulations after adding noise (b), and experimental fcGMP fluorescence traces (c). The reduction in long-lived doubly bound events as compared to singly-bound events in the simulated data is a consequence of the fact that unbinding of either ligand will exit a doubly-bound state.

**Supplementary Fig. 13.**
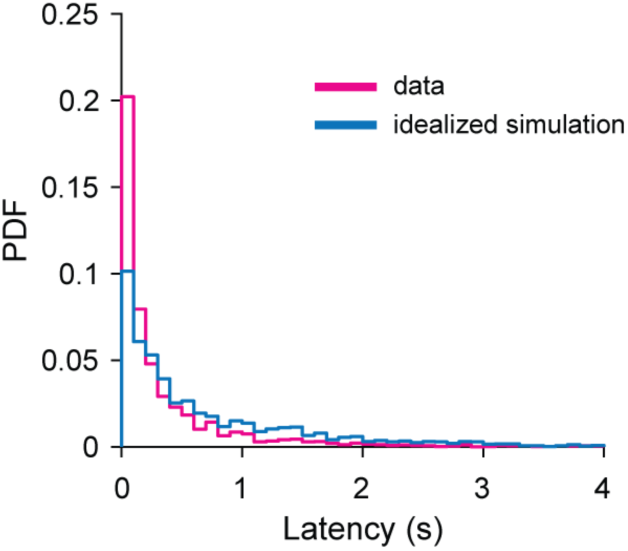
Latency between 1^st^ and 2^nd^ binding events. Distribution of latencies to binding of the second ligand after binding the first ligand. Comparison between our experimental data (magenta) and simulations for four independent sites (blue, see Methods in main text

**Supplementary Fig. 14.**
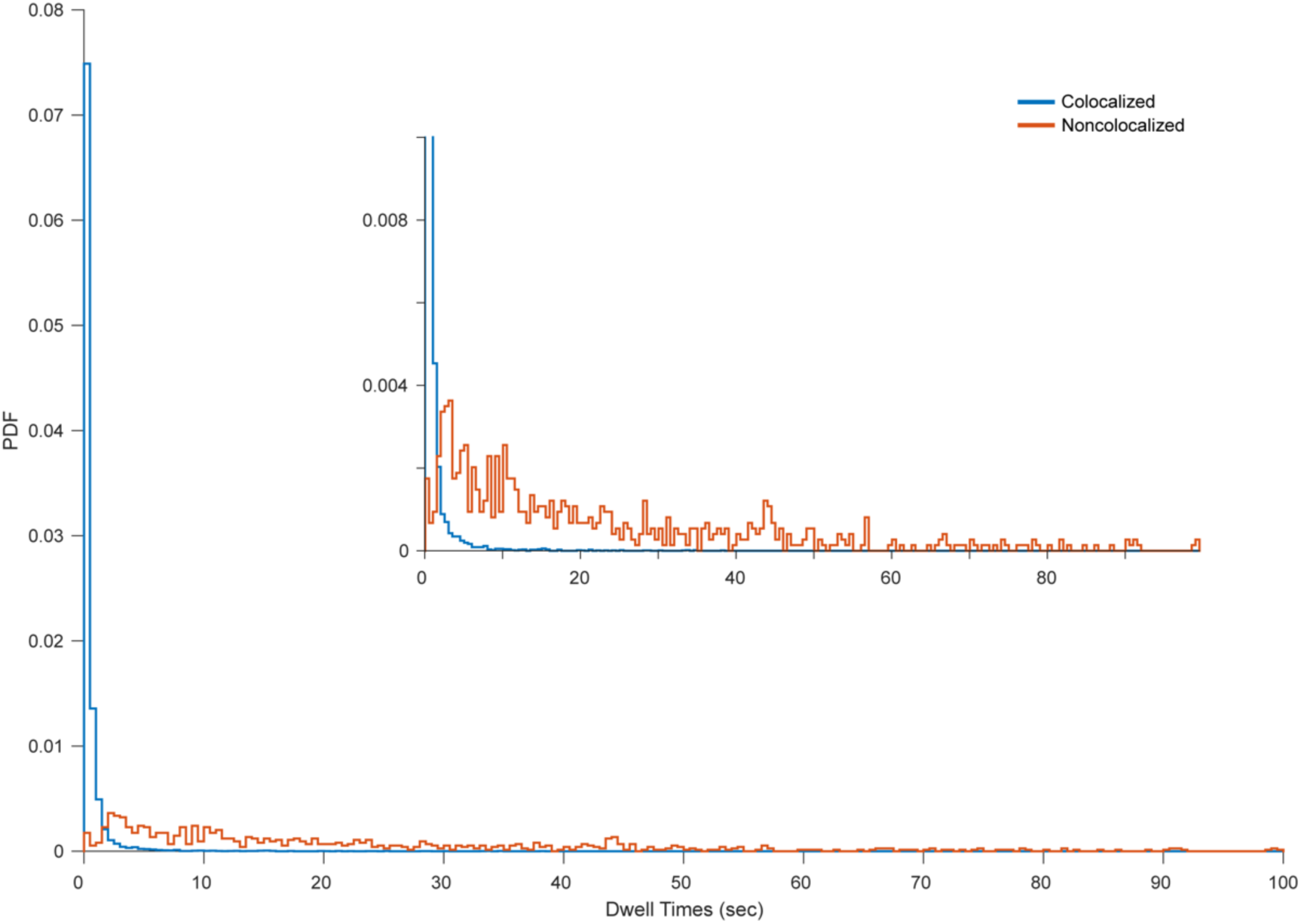
Photobleaching of single fcGMP molecules. Lifetimes of individual fcGMP molecules at noncolocalized spots assumed to largely reflect dye adsorption to the surface and subsequent terminatin by photobleaching. The abscissa is limited to the first 100 seconds for visualization, although lifetimes up to 143 seconds were observed. Lifetimes of colocalized binding events at TAX-4 channels in 30 nM fcGMP are shown for comparison. The mean lifetime for noncolocalized events was 23.4 seconds, an order of magnitude longer than the time constant for the longest duration bound component (Supplementary Table 1).

**Supplementary Table 1.** Time constants (τ) and relative weights (A) for biexponential maximum likelihood fits to bound and unbound dwell time distributions across tested fcGMP concentrations.

| [fcGMP]<br>(nM) | Bound Durations |  |  |  | Unbound Durations |  |  |  |
| --- | --- | --- | --- | --- | --- | --- | --- | --- |
| | $\tau_1$ (s) [95% CI] | $\tau_2$ (s) [95% CI] | A1 [95% CI] | A2 | $\tau_1$ (s) [95% CI] | $\tau_2$ (s) [95% CI] | A1 [95% CI] | A2 |
| 10 | 0.05 [0.043-0.057] | 1.6 [1.5-1.7] | 0.81 [0.78 - 0.83] | 0.19 | 4.2 [3.9-4.5] | 21 [19-23] | 0.76 [0.68 - 0.84] | 0.24 |
| 30 | 0.17 [0.16-0.18] | 1.8 [1.7-1.9] | 0.83 [0.8 - 0.86] | 0.17 | 3.8 [3.6-4] | 12 [11-13] | 0.75 [0.65 - 0.85] | 0.25 |
| 60 | 0.19 [0.17-0.21] | 1.6 [1.4-1.8] | 0.85 [0.79 - 0.91] | 0.15 | 2.1 [1.7-2.5] | 11 [10-12] | 0.85 [0.83 - 0.87] | 0.15 |
| 100 | 0.15 [0.14-0.16] | 2.5 [2.3-2.7] | 0.79 [0.75 - 0.83] | 0.21 | 1.7 [1.5-1.9] | 4.1 [2-6.2] | 0.55 [0.45 - 0.65] | 0.45 |
| 200 | 0.18 [0.17-0.19] | 2.4 [2.3-2.5] | 0.81 [0.78 - 0.84] | 0.19 | 1.3 [1.2-1.4] | 5.7 [4.9-6.5] | 0.82 [0.75 - 0.89] | 0.18 |

**Supplementary Table 2.** Rate constants and relative BIC scores for single site models **(Supplementary Fig. 9)**. Units are μM^-1^s^-1^ for binding transitions (***k*_0_**_→**1**_, ***k*_0_**_→**1**\*∗_, ***k*_0*_**_→**1\***_) and s^-1^ for all other transitions. Rate constants were optimized in QuB for data segments comprised of isolated binding events (no stacked events) across all molecules and concentrations (see Methods in main text). For models M1.D and M1.F one of the rate constants was constrained to enforce microscopic reversibility in the loop. Errors are standard deviations across optimized rate constants for five randomized folds of the data set where each fold contained ∼20% of the data for each concentration.

| Model | $k_{0 \rightarrow 1}$ | $k_{1 \rightarrow 0}$ | $k_{1 \rightarrow 1^*}$ | $k_{1^* \rightarrow 1}$ | $k_{0 \rightarrow 0^*}$ | $k_{0^* \rightarrow 0}$ | $k_{0^* \rightarrow 1^*}$ | $k_{1^* \rightarrow 0^*}$ | $k_{0 \rightarrow 1^*}$ | $k_{1^* \rightarrow 0}$ | $\Delta\text{BIC}$ |
| --- | --- | --- | --- | --- | --- | --- | --- | --- | --- | --- | --- |
| M1.A | 4.2<br>$\pm 0.2$ | 2.3<br>$\pm 0.04$ | — | — | — | — | — | — | — | — | 17175 |
| M1.B | 4.4<br>$\pm 0.2$ | 4.5<br>$\pm 0.1$ | 0.54<br>$\pm 0.06$ | 0.54<br>$\pm 0.05$ | — | — | — | — | — | — | 5412 |
| M1.C | 3.8<br>$\pm 0.1$ | 5.1<br>$\pm 0.2$ | — | — | — | — | — | — | 0.58<br>$\pm 0.07$ | 0.47<br>$\pm 0.04$ | 5412 |
| M1.D | 4.4<br>$\pm 0.2$ | 4.5<br>$\pm 0.1$ | 0.54<br>$\pm 0.06$ | 0.54<br>$\pm 0.05$ | — | — | — | — | 5.7e-8<br>$\pm 0.1\text{e-}8$ | 5.8e-7<br>$\pm 0.1\text{e-}7$ | 5427 |
| M1.E | 7.0<br>$\pm 0.3$ | 4.5<br>$\pm 0.1$ | 0.54<br>$\pm 0.06$ | 0.54<br>$\pm 0.05$ | 0.06<br>$\pm 0.003$ | 0.17<br>$\pm 0.01$ | — | — | — | — | 797 |
| M1.F | 7.4<br>$\pm 0.3$ | 4.7<br>$\pm 0.1$ | 0.25<br>$\pm 0.04$ | 0.27<br>$\pm 0.03$ | 0.03<br>$\pm 0.001$ | 0.05<br>$\pm 0.002$ | 0.74<br>$\pm 0.37$ | 0.31<br>$\pm 0.02$ | — | — | 0 |

**Supplementary Table 3.** Rate constants and relative BIC scores for models of the first two binding steps **(Supplementary Fig. 10)**. Units are μM^-1^s^-1^ for binding transitions (*k*_0→1_, *k*_1→2_, *k*_0*→1*_, *k*_1*→2*_) and s^-1^ for all other transitions. Gray shaded cells indicate constraints, and all other cells were free parameters. All constraints for *k*_1→2_ and *k*_2→1_ reflect the statistical factors associated with independent binding at each site in a tetramer, whereas otherwise the binding steps were allowed to differ (i.e. be cooperative). Constraints for model M2.C transitions *k_i_*_→*i*\*_ and *k_i*_*_→*i*_ reflect the additive effect of ligand binding on the conformational change between states *i* and *i*\* as depicted by the factors *α* and *β* in Supplementary Fig. 10. For all three M2.D models, constraints on transitions not involving ligand binding/unbinding imply that the ratio of the transition rate for the conformational change in doubly liganded states (2 → 2*) to that in singly liganded states (1 → 1*) is given by %, and the factor of 2 is a statistical factor given that either site could undergo the conformational change. Likewise, the ratio of the reverse transition rates for 2* → 2 and 1* → 1 is given by *β*. Constraints for all M2.D models not shown in the table are *k*_1*→2*_ = α*k*_1→2_, *k*_2*→1*_ = (β⁄2) *k*_2→1_, *k*_2*→2**_ = (α⁄2) *k*_2→2*_ and *k*_2**→2*_ = 2*β k*_2*→2_. Rate constants were optimized in QuB for the entire data set across all molecules and concentrations excluding events with more than two bound ligands (see Methods in main text). Errors are standard deviations across optimized rate constants for five randomized folds of the data set where each fold contained ∼20% of the data for each concentration.

| Model | $k_{0 \rightarrow 1}$ | $k_{1 \rightarrow 0}$ | $k_{1 \rightarrow 2}$ | $k_{2 \rightarrow 1}$ | $k_{1 \rightarrow 2^\dagger}$ | $k_{2^\dagger \rightarrow 1}$ | $k_{0 \rightarrow 0^*}$ | $k_{0^* \rightarrow 0}$ | $k_{1 \rightarrow 1^*}$ | $k_{1^* \rightarrow 1}$ | $k_{2 \rightarrow 2^*}$ | $k_{2^* \rightarrow 2}$ | $\alpha$ | $\beta$ | $\Delta\text{BIC}$ |
| --- | --- | --- | --- | --- | --- | --- | --- | --- | --- | --- | --- | --- | --- | --- | --- |
| M2.A<sub>i</sub> | $4.4 \pm 0.1$ | $1.6 \pm 0.04$ | $\frac{3}{4} k_{0 \rightarrow 1}$ | $2 k_{1 \rightarrow 0}$ | — | — | — | — | — | — | — | — | — | — | 27209 |
| M2.A<sub>c</sub> | $4.4 \pm 0.1$ | $1.8 \pm 0.1$ | $3.4 \pm 0.1$ | $1.6 \pm 0.1$ | — | — | — | — | — | — | — | — | — | — | 24584 |
| M2.B | $4.4 \pm 0.1$ | $1.9 \pm 0.1$ | $3.0 \pm 0.1$ | $4.2 \pm 0.2$ | $0.50 \pm 0.03$ | $0.30 \pm 0.02$ | — | — | — | — | — | — | — | — | 22191 |
| M2.C<sub>i</sub> | $8.0 \pm 0.2$ | $3.6 \pm 0.1$ | $\frac{3}{4} k_{0 \rightarrow 1}$ | $2 k_{1 \rightarrow 0}$ | — | — | $0.11 \pm 0.02$ | $0.24 \pm 0.03$ | $0.51 \pm 0.04$ | $0.55 \pm 0.03$ | $2.1 \pm 0.1$ | $0.84 \pm 0.02$ | — | — | 1338 |
| M2.C<sub>c</sub> | $7.0 \pm 0.1$ | $4.2 \pm 0.1$ | $8.5 \pm 0.3$ | $3.7 \pm 0.1$ | — | — | $0.06 \pm 0.01$ | $0.18 \pm 0.02$ | $0.77 \pm 0.05$ | $0.68 \pm 0.03$ | $0.50 \pm 0.03$ | $0.40 \pm 0.03$ | — | — | 0 |
| M2.D | $8.0 \pm 0.2$ | $3.8 \pm 0.1$ | $\frac{3}{4} k_{0 \rightarrow 1}$ | $2 k_{1 \rightarrow 0}$ | — | — | $\frac{1}{\alpha} k_{1 \rightarrow 1^*}$ | $\frac{1}{\beta} k_{1^* \rightarrow 1}$ | $0.01 \pm 0.001$ | $0.01 \pm 0.001$ | $\alpha k_{1 \rightarrow 1^*}$ | $\beta k_{1^* \rightarrow 1}$ | $0.18 \pm 0.005$ | $0.11 \pm 0.003$ | 1537 |
| M2.E<sub>ii</sub> | $4.6 \pm 0.1$ | $3.9 \pm 0.1$ | $\frac{3}{4} k_{0 \rightarrow 1}$ | $2 k_{1 \rightarrow 0}$ | — | — | — | — | $0.46 \pm 0.04$ | $0.34 \pm 0.02$ | $2\alpha k_{1 \rightarrow 1^*}$ | $\beta k_{1^* \rightarrow 1}$ | 1 | 1 | 4550 |
| M2.E<sub>ic</sub> | $4.8 \pm 0.1$ | $4.1 \pm 0.1$ | $\frac{3}{4} k_{0 \rightarrow 1}$ | $2 k_{1 \rightarrow 0}$ | — | — | — | — | $0.53 \pm 0.04$ | $0.44 \pm 0.02$ | $2\alpha k_{1 \rightarrow 1^*}$ | $\beta k_{1^* \rightarrow 1}$ | $0.75 \pm 0.04$ | $0.49 \pm 0.02$ | 3834 |
| M2.E<sub>ci</sub> | $4.7 \pm 0.1$ | $4.3 \pm 0.1$ | $3.5 \pm 0.1$ | $4.1 \pm 0.2$ | — | — | — | — | $0.44 \pm 0.04$ | $0.34 \pm 0.02$ | $2\alpha k_{1 \rightarrow 1^*}$ | $\beta k_{1^* \rightarrow 1}$ | 1 | 1 | 3236 |

